# Increased variation in sex organ positions can increase reciprocity and pollination success in heterostylous plant populations

**DOI:** 10.1101/2020.02.04.934760

**Authors:** Shatarupa Ganguly, P.M. Shreenidhi, Deepak Barua

## Abstract

**Aims:** The reciprocal position of sexual organs in complementary floral morphs is central to our understanding of heterostyly. Reciprocity indices are used to quantify the spatial match between complementary sex organs, but previous indices fail to appropriately account for variation in sex organ positions among individuals in a population. The objective of this study was to examine how reciprocity and consequently reproductive success change with an increase in intra-population variation in sex organ heights. To this end, we formulated a reciprocity index that incorporates variation in sex organ positions among individuals in a population and asked if estimates of reciprocity can predict reproductive success in naturally occurring heterostylous populations.

**Methods:** We developed a reciprocity index that assumed pollen transfer success equalled one for a perfectly matched stigma-anther pair, and decreased to zero with increasing mismatch. Reciprocity was quantified as the average pollen transfer success for all pair-wise combinations of complementary sex organs in the population. We examined the relationship between intra-population variation and reciprocity using simulated populations that varied in the distribution of sex organ positions, and with empirical data from natural populations. We compared previously proposed indices using the simulated and natural populations, and for a subset of natural populations we tested the ability of the indices to predict reproductive success.

**Important Findings:** In both simulated and natural populations we observed that when differences between mean anther and stigma heights of complementary morphs are small, increasing intra-population variation in heights resulted in a monotonous decrease in reciprocity. However, when differences between mean complementary anther and stigma heights are larger, reciprocity increased, reached a peak, and then decreased with increasing variation. Previous indices failed to capture this behaviour and were largely insensitive to variation or differences in mean complementary sex organ heights. Seed set was consistently positively related to reciprocity for our index, and for two of the four previous indices. These results highlight the importance of incorporating intra-population variation in sex organ dimensions in quantifying reciprocity, and challenge the current understanding that increasing variation will always decrease reciprocity in heterostylous populations. These results may help explain why heterostylous systems exhibit, and tolerate high amounts of intra-population variation in sex organ heights.

## INTRODUCTION

The reciprocal arrangement of anthers and stigmas in complementary floral morphs is central to our understanding of the function and evolution of heterostyly (Darwin, 1877; Lloyd and Webb, 1992). This reciprocal arrangement ensures that pollen from anthers of a morph is deposited on the pollinator’s body to match the position of the stigmas in the complementary morph, and deviation from this reduces the precision of pollen transfer (Stone and Thomson, 1994; Armbruster *et al*., 2009; Keller *et al*., 2012; Brys and Jacquemyn, 2015; Liu *et al*., 2016). This spatial match between anthers and stigmas is widely quantified using reciprocity indices, and used to estimate potential pollen transfer and reproductive success in heterostylous populations (Richards and Koptur, 1993; Eckert and Barrett, 1994; Sanchez *et al*., 2008; Sosenski *et al*., 2010; Keller *et al*., 2012; Sánchez *et al*., 2013; Armbruster *et al*., 2017). The commonly used reciprocity indices incorporate mean anther and stigma positions but fail to appropriately account for intra-population variation in organ heights.

Distylous species have long-, and short-styled individuals with anthers and stigmas at two distinct levels (Fig. 1 A). Legitimate pollen transfer takes place between anthers and stigmas of complementary morphs at the same level. Differences in the height of sex organs at the same level in complementary morphs results in a decrease in reciprocity (Richards and Koptur, 1993). In addition to differences in mean complementary sex organ heights, variation in heights among individuals in the population can also result in a decrease in reciprocity. Such Intra-population variation in sex organ position has been widely reported, and is likely common in heterostylous plants (Eckert and Barrett, 1994; Pailler and Thompson, 1997; Faivre and McDade, 2001; Sanchez *et al*., 2008; Ferrero, 2009). The extent of such variation can be substantial (Wolff and Liede-Schumann, 2007; Brys *et al*., 2008; Chen, 2009; Machado *et al*., 2010), and can reduce pollen transfer between morphs, and reproductive success in the population (Ferrero, Chapela, *et al*., 2011; Keller *et al*., 2012; Brys and Jacquemyn, 2015; Haller *et al*., 2014).

**Figure 1:**
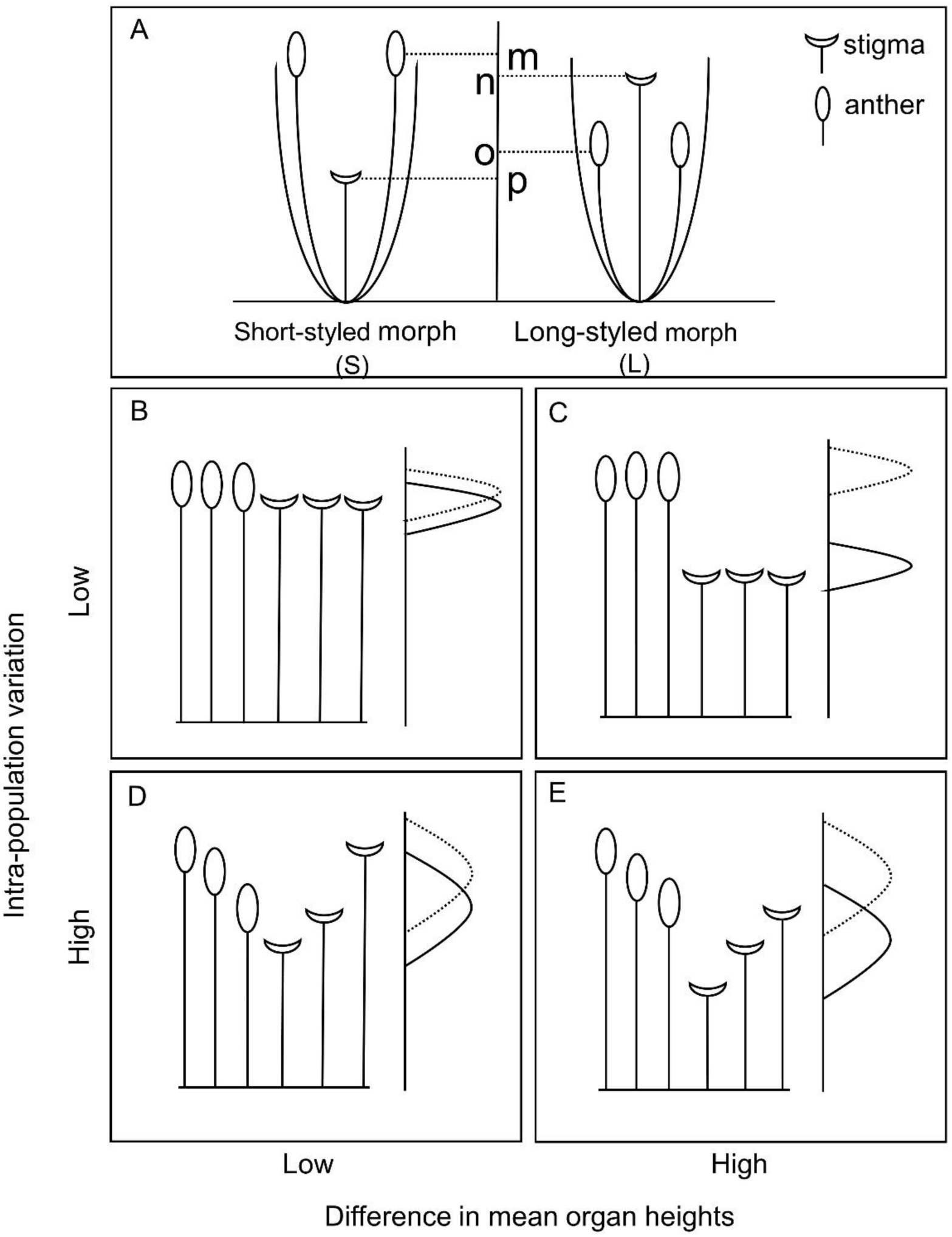
A) Schematic for a long-styled and a short-styled distylous flower. The measurements from the base of the flower to points m and o – anther heights of short-styled morph and long-styled morph respectively; and, to points n and p – stigma heights of long-styled morph and short-styled morph respectively. Thus, m-n and o-p represent difference in mean organ heights for the high and low sexual organ position respectively. B-E) Pictorial representation of low and high difference in mean sexual organ heights and intra-population variation on reciprocity. The right side of each panel shows the vertical distribution of complementary stigma (bold line) and anther (dotted line) heights and the left side of each panel shows a pictorial representation of 3 individuals each of complementary morphs at one level of sexual organ position of heterostylous plants.

Early indices developed to quantify reciprocity focused on mean organ heights, and neglected intra-population variation (Richards and Koptur, 1993). Eckert and Barrett (1994) proposed a precision index based on the coefficient of variation in sex organ heights to account for intra-population variation, but provided no explanation for how to combine this measure with the reciprocity index. Later indices recognised the importance of intra-population variation in organ positions and incorporate estimates of variation in quantifying reciprocity (Lau and Bosque, 2003; Sanchez *et al*., 2008; Sánchez *et al*., 2013; Armbruster *et al*., 2017). However, the manner in which this was done for the different indices differed, little justification was provided for the mathematical operations used, and the interpretation of these quantitative estimates of reciprocity remained problematic.

The index proposed by (Lau and Bosque, 2003) calculates reciprocity as the percentage overlap between the frequency distributions of complementary anther and stigma heights of a population. However, these estimates of reciprocity are sensitive to the choice of bin sizes used for the frequency distributions of the sex organ heights. This choice of bin size is arbitrary with no apparent biological significance. The calculation of Sanchez’s index requires multiplication of the spatial reciprocity estimates with the standard deviation in organ heights, but the biological justification for this is unclear (Sanchez *et al*., 2008; Sánchez *et al*., 2013).

Higher intra-population variation in traits is often associated with decreased fitness due to deviation from an optimal trait value. However, higher intra-population variation can be beneficial under certain circumstances (Crean and Marshall, 2009), and this has also been documented for sex-organ positions in plants (Dai *et al*., 2016). In heterostylous plants, the potential matches for successful pollen transfer for an individual with anthers of a particular height is represented by the range of stigmas heights in individuals of the complementary morph in the population. Thus, reciprocity depends on the mean complementary stigma height, but also on the distribution of stigma heights in the population. When the mean heights of the complementary sex organs are perfectly matched, increasing intra-population variation in sex organ height, will result in a decrease in reciprocity (Fig. 1 B, D). However, when there is a mismatch in mean complementary sex organ heights, low intra-population variation in sex organ heights will actually result in no overlap in complementary anthers and stigmas, and reciprocity will be zero (Fig. 1 C). In such situations, increasing intra-population variation in sex organ heights, will ultimately result in an increase in reciprocity as the tallest stigmas will overlap with the shortest anthers (Fig. 1 E).

The reciprocity index developed in this study (henceforth current index) quantifies mismatch in vertical heights of complementary anthers and stigmas in distylous plant populations. For every combination of anther and complementary stigma, we assign a pollen transfer success value that is inversely related to mismatch. Thus, pollen transfer success equals one for a perfect match and decreases with increasing mismatch between the stigma and anther. We quantify reciprocity as the average pollen transfer success for all pair-wise combinations of anthers and stigmas in the population. One important difference from previous indices in that we convert the spatial match in anther-stigma pairs to a pollen transfer success between sex organs, a measure that reciprocity inherently tries to capture.

We used the current index to examine how reciprocity changes with increasing intra-population variation in sex organ heights. For this we used simulated plant populations that varied in mean sex organ heights, and intra-population variation in heights. We extracted data on sex organ heights for heterostylous populations from previously published studies to understand the biologically relevant ranges of mean and intra-population variation in anther and stigma positions, and used this information to determine the ranges for these parameters in our simulated population. Using these simulated populations, we also compared the estimates of reciprocity from the current index to the previously proposed indices. In parallel, we used the data extracted from previous studies that represented the combinations of mean and intra-population variation in sex organ positions in natural populations, to understand how reciprocity changes with increasing intra-population variation in organ heights, and to compare the current index to the previously proposed ones. While reciprocity indices are commonly used to understand potential reproductive success in heterostylous plants, the relationship between proposed indices and measures of reproductive success have rarely been tested (Jacquemyn *et al*., 2018; Brys and Jacquemyn, 2019). To do this, we examined the relationship between the current and previous indices to reproductive success using data on fruit and seed set in heterostylous plants.

## METHODS

### Sex organ positions in natural heterostylous populations

We extracted data on sex organ heights in distylous species to understand: a) the relative position of the lower-, and higher-level sex organs; b) differences in mean complementary sex organ positions; and, c) intra-population variation in sex organ heights. This was used to determine the relative position of sex organ heights in the simulated flowers, and the range of sex organ heights and intra-population variation that we explored in our simulated populations. We used the keywords distyly/distylous or heterostyly/heterostylous to identify papers that were screened to determine if they contained individual-level information for sex organ heights. We only used studies that reported data for more than 10 individuals per morph. We considered data for species and populations independently, i.e. different populations of the same species were treated as independent data points for further analysis. Plot Digitizer (Joseph A. Huwaldt, http://plotdigitizer.sourceforge.net/) was used to extract information from figures or graphs. The difference in mean complementary sex organ position for each level, and standard deviation for sex organ heights were calculated for each morph. Since, flower size varied between species, the values of mean sex organ position and intra-population variation for all four sex organs were standardized using the grand mean of the mean anther and stigma heights of the high level. The mean anther and stigma height of the high level was assumed to be a proxy for flower size as information for the latter was not always available. When available we also extracted information from these studies on anther length for the study species, and fruit set and/or seed set for the study populations.

### Calculation of the reciprocity index

To quantify reciprocity we calculated the mismatch for every combination of complementary anther and stigma in the population. Mismatch, *MM* was defined as the modulus of the difference in vertical position of the anther and stigma, and this was calculated separately for the lower and higher organ levels.

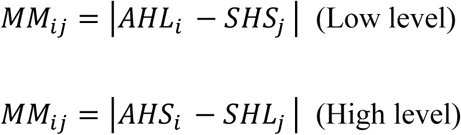

Where, *AHL* is the anther height of the long-styled morph; *SHS* is the stigma height of the short-styled morph; *AHS* is the Anther height of the short-styled morph; *SHL* is the stigma height of the long-styled morph for the *i*th and *j*th individual of the long-, or the short-styled morph, respectively (Fig. 1 A).

We assumed that pollen transfer success decreased with increasing mismatch. Given the lack of empirical information regarding how anther and stigma position translate to pollen transfer between complementary sex organs, we initially used a concave-shaped decreasing function to convert each value of mismatch to a pollen transfer success value, *R_ij_*. This assumed that the consequences of mismatch were more severe for changes in pollen transfer initially, and less severe later as the mismatch increased (Washitan *et al*., 1994; Deschepper *et al*., 2018).

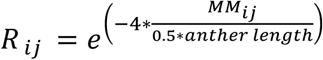

Where *R_ij_* is pollen transfer success, and M*_ij_* the mismatch between the *i*th anther and *j*th stigma of complementary morphs. Values for *R_ij_* ranged from one for a perfectly matched stigma-anther pair, to zero when the mismatch is greater than half of the anther length (Ferrero, Chapela, *et al*., 2011). In a later section we examined the consequences of relaxing these assumptions: using different decreasing functions; as well as different critical values of mismatch at which pollen transfer success becomes zero.

The average pollen transfer success for all possible combinations of anthers and stigmas was calculated independently for both high-, and low-level organs in the population.

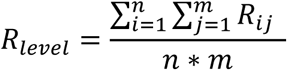

Here level refers to either high-, or low-level of a distylous flower and *n* and *m* are the numbers of individuals of the short-styled and long-styled morph, respectively.

### Explorations with simulated populations

We used simulated populations of heterostylous plants to examine how our measure of reciprocity changed with mean sex organ heights and increasing intra-population variation. The relative positions for the higher-level anther and stigma in the simulated individuals were set at 100, and the low-level was determined based on the empirical data obtained from the naturally occurring heterostylous populations. For the simulations, to ensure that the relative positions of the sex organs in the two levels stayed the same, when we increased the difference in mean complementary sex organ heights, we distributed the difference equally amongst the mean anther and stigma heights of the complementary sex organs at that level.

We used normal distributions of anther and stigma heights for 200 individuals with 100 individuals of each morph. To examine how sex organ height affects reciprocity, we estimated reciprocity across a range of intra-population variation in heights, and for a range of mean height differences between complementary sex organs. The lower and upper bounds of the range for both intra-population variation and height differences were determined from the empirical data from naturally occurring heterostylous populations extracted from previously published studies. We assumed that the intra-population variation and difference in mean complementary sex organs heights were similar for both high and low levels. We used the average value of the estimated reciprocity from 100 simulation runs for each combination of mean sex organ heights and intra-population variation examined.

### Comparison of pollen transfer success functions and different critical values of mismatch

To understand if the results were robust to different pollen transfer success functions, we compared three decreasing functions: (a) convex; (b) linear; and (c) concave [Supplementary information Fig. S1]. Additionally, we also examined a uniform function where the pollen transfer success was independent of mismatch when mismatch was less than half anther length, and was zero when mismatch was greater than half anther length. These functions represent a gradient of increasing penalty with respect to pollen transfer success with increasing mismatch in the following order: uniform < convex < linear < concave. We also examined how the critical value of mismatch at which pollen transfer success becomes zero affects reciprocity. For this, the critical value was set at 5%, 10%, 15% and 20% of the average height of the higher level which is used as a measure of flower size.

### Comparison of indices with simulated populations

Using the simulated populations, the current index was compared to four other commonly used reciprocity indices: a) (Richards and Koptur, 1993); b) (Lau and Bosque, 2003); c) (Sánchez *et al*., 2013); and d) (Armbruster *et al*., 2017). Richards & Koptur’s index was modified as suggested by (Sanchez *et al*., 2008) to incorporate intra-population variation. We used a subset of difference in mean complementary sex organ heights of the simulated populations representing high and low values of the explored range, to compare the indices. To ease comparison with the other indices, the reciprocity values for Armbruster *et al*. (2017) were scaled and flipped to range between zero and one by first dividing each value by the maximum value of the index obtained across all difference in mean complementary sex organ heights and then subtracting from one.

### Explorations with naturally occurring populations

We used the individual-level empirical data obtained from naturally occurring heterostylous populations to examine how our measure of reciprocity changed with mean sex organ heights and increasing intra-population variation. As with the simulated populations, we used half of the mean anther length for the critical value of mismatch at which pollen transfer success becomes zero. Anther length information was extracted from the same study from which we extracted data on organ heights. When not unavailable, we used the mean anther length for the genus or family of the species.

To examine how increasing intra-population variation in organ heights affected reciprocity we pooled species in four categories based in differences in mean complementary sex organ heights. The four categories expressed as the percentage of the grand mean of mean high-level sex organs were: 0 - 5%; 5 - 10%; 10 - 15%; and, > 15%. Within each of these categories we examined the relationship between reciprocity and intra-population variation in sex organ heights for the high and low level.

### Comparison of indices using empirical data from naturally occurring populations

We used the empirical data from naturally occurring heterostylous populations to compare estimates of reciprocity using the current and previous indices. This allowed us to compare the indices for biologically relevant combinations of sex organ heights and intra-population variation. The reciprocity for each population was calculated separately for the higher and lower sex organ level, except for the Sanchez *et al*. (2013) index that estimates reciprocity for the population as a whole. For all comparisons with the Sanchez *et al*. (2013) index, we use the average difference in mean complementary sex organ heights of both organ levels and average intra-population variation of all four sex organ heights. Values for Armbruster *et al*. (2017) were standardized by dividing with the grand mean of all four mean sex organ heights for that species (Armbruster *et al*. 2017). This value was then scaled and flipped to range between zero and one as described before. Armbruster’s original index ranges from zero for perfect reciprocity to infinity. Standardizing and flipping this helped with comparison with the other indices. The reciprocity values for the high and low levels were categorised based on the difference in mean complementary sex organ heights as mentioned in the previous section.

### Do reciprocity indices predict reproductive success?

Finally, we examined the relationship between estimates of reciprocity for the naturally occurring populations and reproductive success for a subset of studies where information on fruit or seed set for the populations was available. These values of fruit or seed set data expressed as percentage values per flower, or per ovule per flower, respectively, were used a measure of reproductive success. Spearman’s rank correlation coefficients were used to examine the relationship between the reciprocity estimates from the current and previous indices, and fruit or seed set for the upper and lower sex-organ levels separately. For eight populations, fruit set for the short morph was zero, i.e. they were likely functionally dioecious, and these populations were excluded from the analysis.

## RESULTS

### Sex organ positions in natural heterostylous populations

The literature search resulted in the identification of 70 publications with data for 211 populations of 114 species that were used to extract data for sex organ heights [Supplementary Information: Table S1; Supplementary references].

The mean sex organ positions and intra-population variation are presented as a percentage of the grand mean of mean sex organ heights for the higher level for each of the 211 populations of the 114 species. The relative position of the lower sex organ was at approximately 60% of the height of the high level. For the long-, and short-styled morphs, the mean and standard deviation of the position of the lower sex organ level was 60.58% ± 13.95, and 63.91% ± 11.92 respectively. The difference in mean complementary sex organ heights varied widely, and the 95% confidence limits ranged from 0.67 % to 31.99 % (mean ± sd = 10.30 ± 9.01%) for the higher organ level, and from 0.57% to 29.44% (mean ± sd = 9.62 ± 8.68%) for lower organ level (Fig. 2). Intra-population variation in sex organ heights also varied widely and the 95% confidence limits ranged from 4.14% to 16.10% (mean ± sd = 9.65 ± 3.52%) for stigma height, and from 2.93% to 11.96% (mean ± sd = 6.91 ± 3.06%) for anther height of the long-styled morph. The 95% confidence limits for intra-population variation in the short-styled morph ranged from 2.97% to 13.00% (mean ± sd = 7.11 ± 2.95%) for stigma height, and from 3.92% to 15.61% (mean ± sd = 9.07 ± 3.71%) for anther height (Fig. 2). Relative anther length was approximately 20% (mean ± sd = 18.21 ± 8.05%, *n* = 89 populations/ 53 species) of the mean height of the higher-level organs.

**Figure 2:**
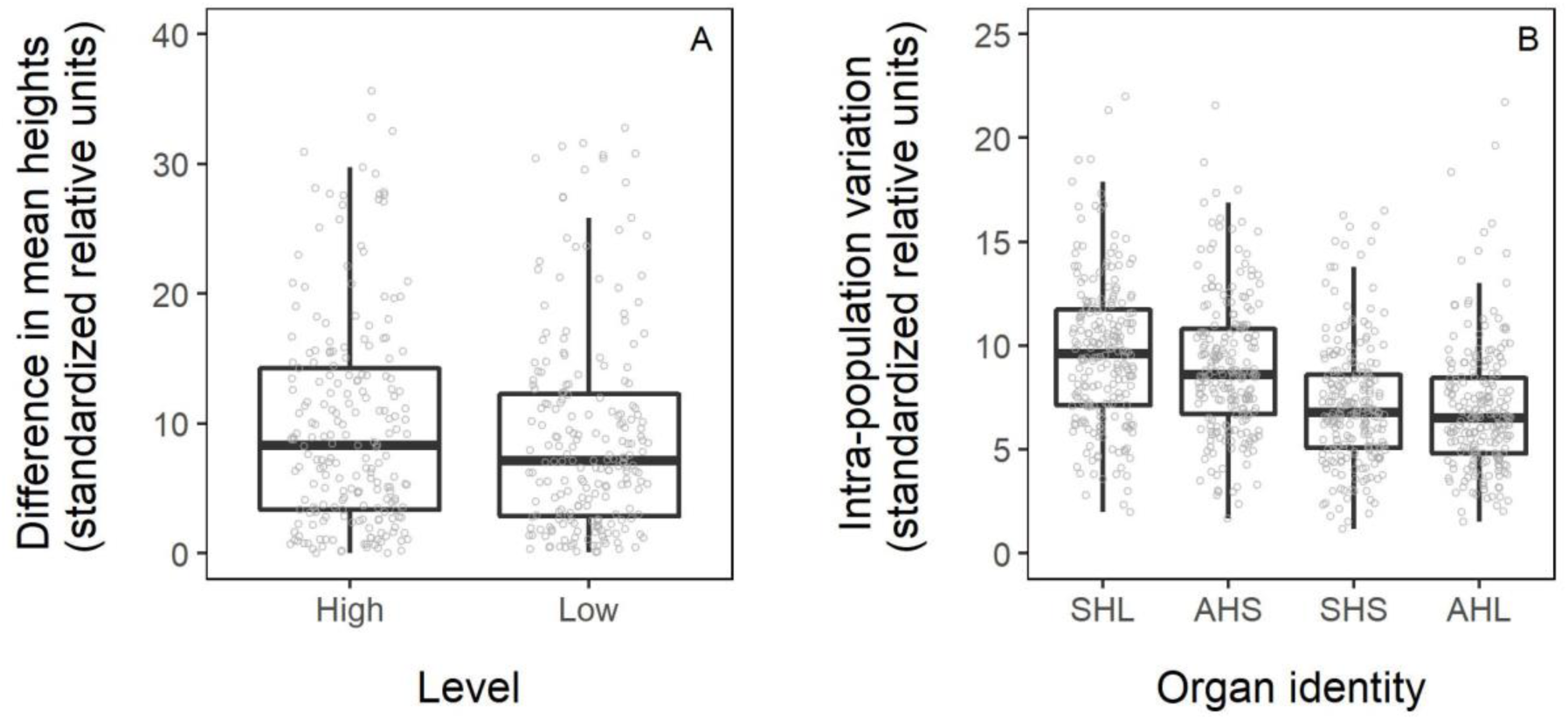
Distribution of difference in mean complementary sex organ heights and intra-population variation extracted from literature: A) Difference in mean complementary sex organ heights for the high and low levels; B) Intra-population variation for the four sex organs. SHL and AHL refer to stigma and anther height of long-styled morph, and SHS and AHS refer to stigma and anther height of short-styled morph (*n* = 211 populations of 114 species).

We used the observed range of sex organ dimensions and intra-population variation to determine the position of the lower sex organs, and the ranges of intra-population variation in sex organ heights and height difference in mean complementary sex organs in the explorations with the simulated populations. The lower sex organ position was fixed at 60% of the higher sex organ positions in our simulated flowers. We selected the range of intra-population variation in sex organ heights and height difference in mean complementary sex organs to explore as 0% - 24%, and 0% - 22% respectively.

### Reciprocity as a function of intra-population variation and difference in mean complementary sex organ heights in simulated populations

For the higher organ level, when differences between mean complementary anther and stigma was zero, the index showed a monotonous decrease in reciprocity with an increase in intra-population variation (Fig. 3 inset). However, when difference in mean complementary sex organ heights was 3%, or greater, of mean height of the high level organs, reciprocity was initially low or zero at low intra-population variation, but increased as intra-population variation increased, reached a peak and then decreased (Fig. 3). The value of intra-population variation at which peak reciprocity was observed increased with increasing differences in mean complementary sex organ heights. The peak value of reciprocity for a given value of intra-population variation, decreased with an increase in difference in mean height of reciprocal sex organs and this decrease is non-linear. Hence, as the difference in mean heights of complementary sex organs increased the difference in peak reciprocity between them decreased. The results for the lower sex organ level were identical. The peak value of the reciprocity was observed at values within the range of difference in mean sex organ heights and intra-population variation as reported in naturally occurring heterostylous plant populations.

**Figure 3:**
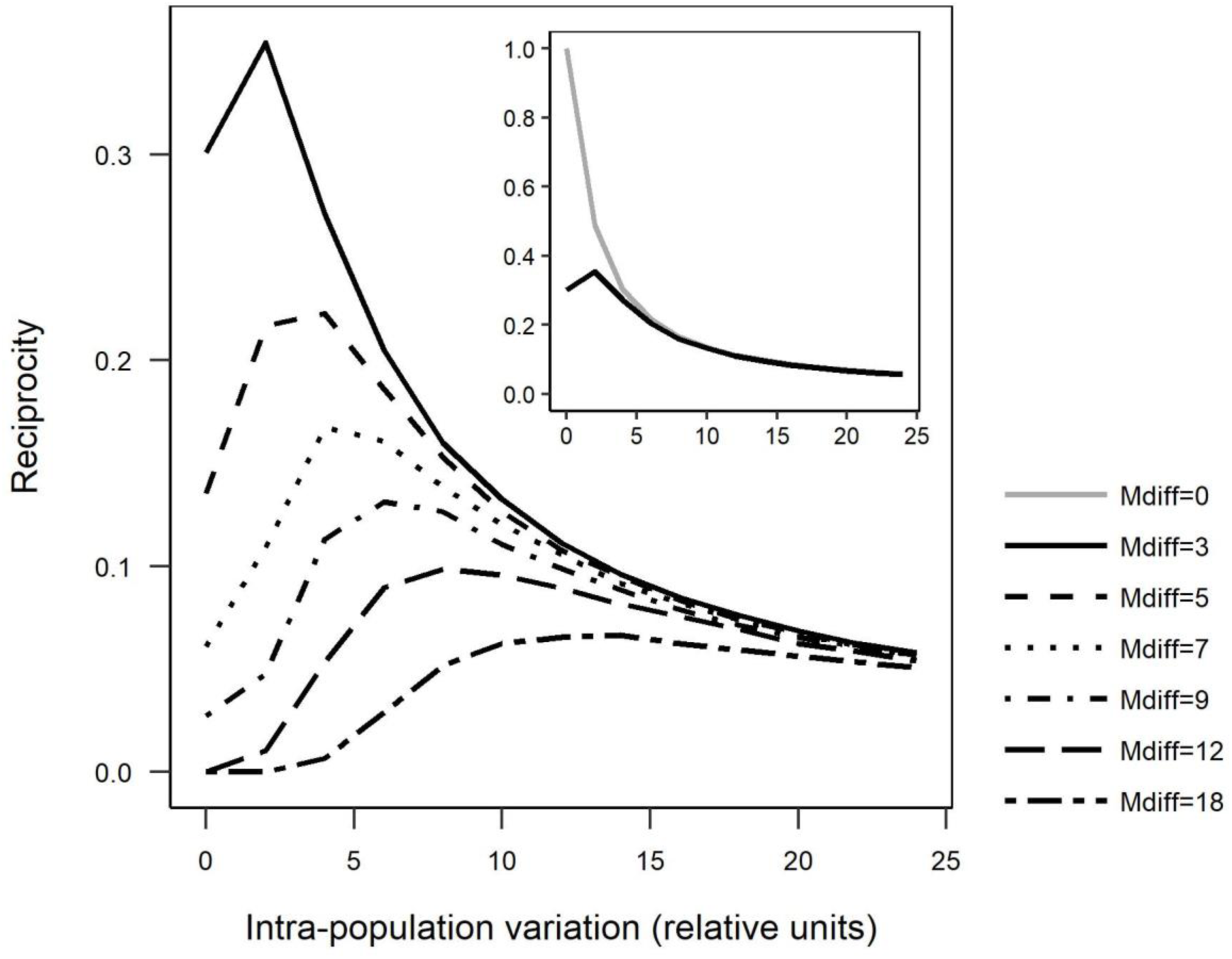
Reciprocity estimates for the simulated populations using the current index as a function of increasing intra-population variation in anther and stigma heights, for a range of values for difference in mean sex organ heights (Mdiff; shown by the different curves). The inset shows reciprocity estimates for lower values of difference in mean sex organ heights (Mdiff = 0, Mdiff = 3)

### Comparison of pollen transfer success functions and different critical values of mismatch

How estimates of reciprocity changed with intra-population variation in sex organ heights and difference in mean complementary sex organ heights were qualitatively similar for the uniform, linear and the convex functions for pollen transfer success examined (Fig. 4). The absolute values of reciprocity differed, and was highest for the uniform function and lowest for the concave function. Changes in the critical value of mismatch at which pollen transfer became zero gave similar results for how reciprocity changed with intrapopulation variation [Supplementary Information: Fig. S2]. However, the absolute values of reciprocity were higher as the critical value increased.

**Figure 4:**
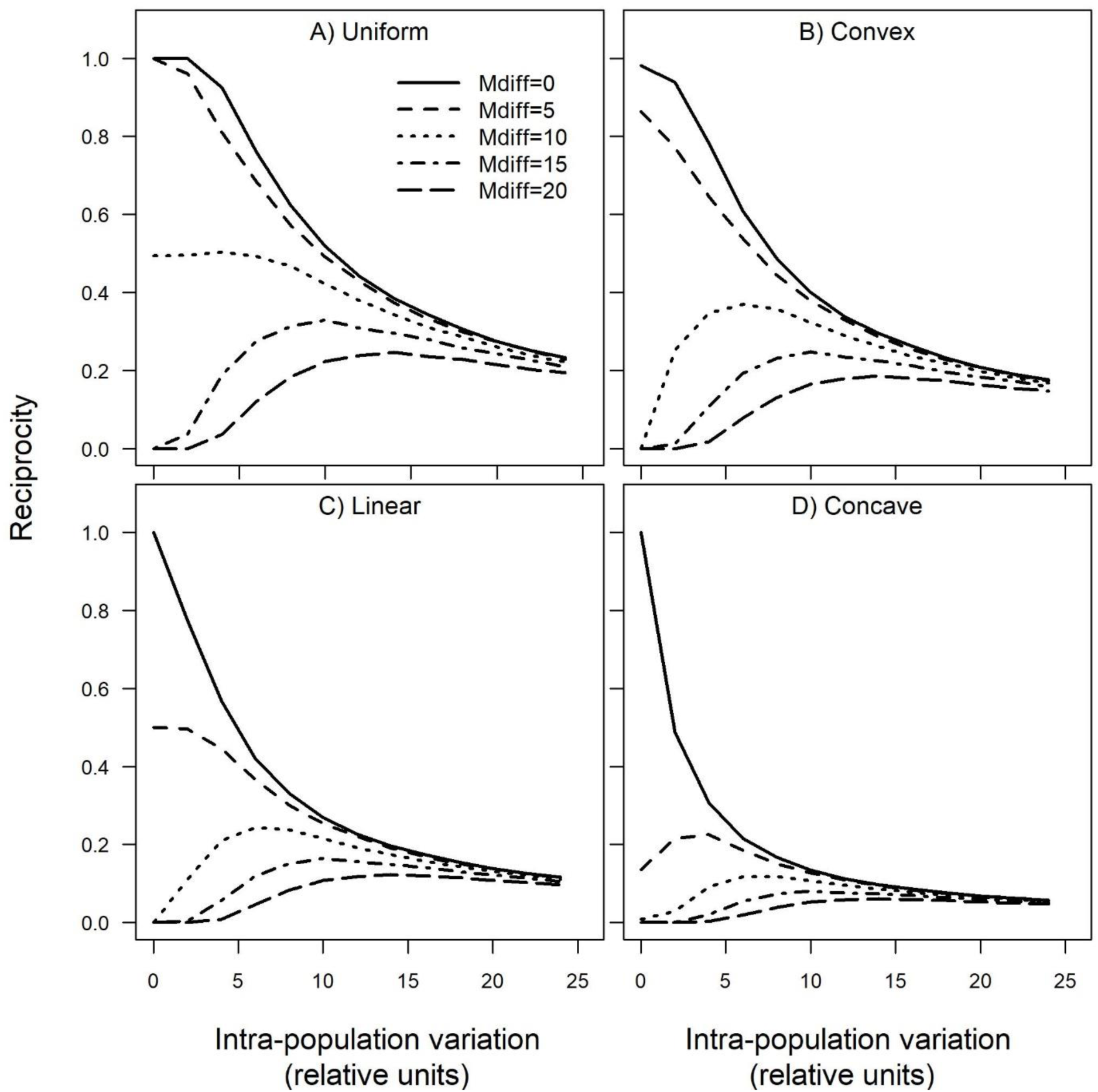
Comparison of the pollen transfer success functions: A) uniform; B) convex; C) linearly; and, D) concave. Reciprocity estimates for the simulated populations as a function of increasing intra-population variation in anther and stigma heights, for a range of values for difference in mean sex organ heights (Mdiff; shown by the different curves).

### Comparison of indices with simulated populations

We observed stark differences in how estimates of reciprocity from the different indices changed with intra-population variation in sex organ heights, and difference in mean complementary sex organs (Fig. 5). When the difference in mean complementary sex organ heights is high (depicted by Fig. 1 C), we expected reciprocity to be low. Additionally, we expected that increases in intra-population variation should result in an increase in reciprocity when difference in mean complementary organs heights were high (compare Fig. 1 C and 1 E). Both of the above were seen for the current index (Fig. 3) and for Lau and Bosque’s index (Fig. 5 B), but not the other three indices. However, unlike the current index, reciprocity estimated by using Lau and Bosque’s index continued to increase and ultimately saturated with increasing intra-population variation (Fig. 5 B).

**Figure 5:**
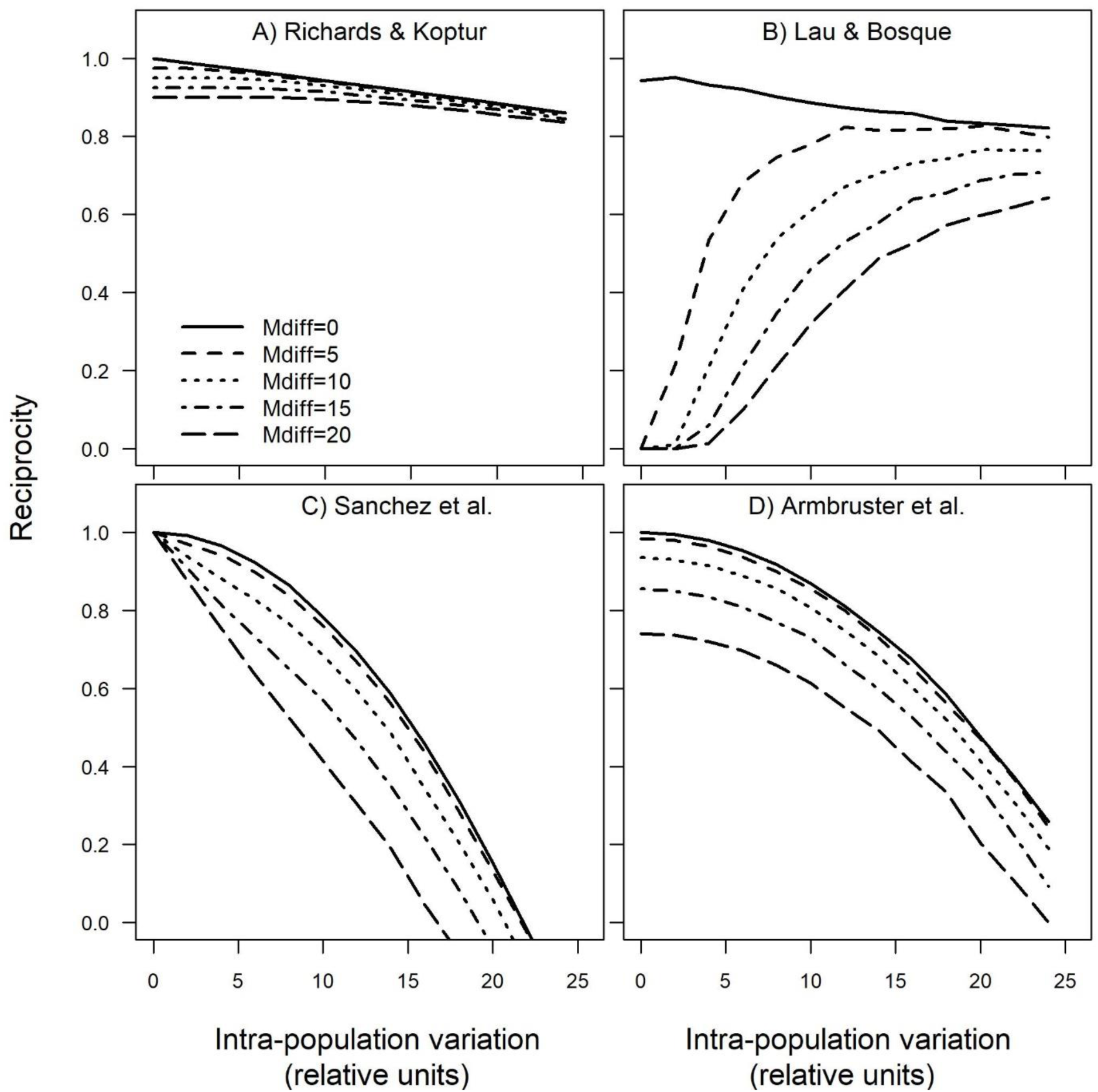
Comparison of the previously proposed indices: A) Richards and Koptur (modified as in Sanchez *et al*. (2008)); B) Lau and Bosque (2003); C) Sanchez *et al*. (2013); and, D) Armbruster *et al*. (2017). Reciprocity estimates for the simulated populations as a function of increasing intra-population variation in anther and stigma heights, for a range of differences in mean sex organ heights (Mdiff; shown by the different curves). Estimates for Armbruster *et al*. (2017) have been scaled to range between zero (minimum) and one (maximum).

The reciprocity indices proposed by Sanchez, Armbruster and the modified Richards & Koptur’s index decreased monotonously with increasing intra-population variation (Fig. 5 A, C & D). The index proposed by Sanchez *et al*. (2013) gave negative values at very high values of difference in mean complementary sex organ heights and intra-population variation (Fig. 5 C). The modified Richards & Koptur’s index was insensitive to changes in both intra-population variation and differences in mean complementary sex organ heights, and resulted in high values of reciprocity throughout the range of intra-population and organ heights examined (Fig. 5 A). Surprisingly, with the exception of Lau & Bosque (2003), all other previous indices were largely insensitive to changes in difference in mean complementary sex organ heights.

### Reciprocity as a function of intra-population variation and difference in mean complementary sex organ heights in naturally occurring populations

In examining the relationship between intra-population variation and reciprocity in naturally occurring populations we saw similar results as with the simulated populations. The current index showed a monotonous decline in reciprocity with increase in intra-population variation when the difference in mean complementary sex organ heights is low (Fig. 6, Mdiff = 0-5). But, when the difference in mean complementary sex organ heights is high (Fig. 6, Mdiff > 5) reciprocity increases with an increase in intra-population variation reaches a peak and then decreases.

**Figure 6:**
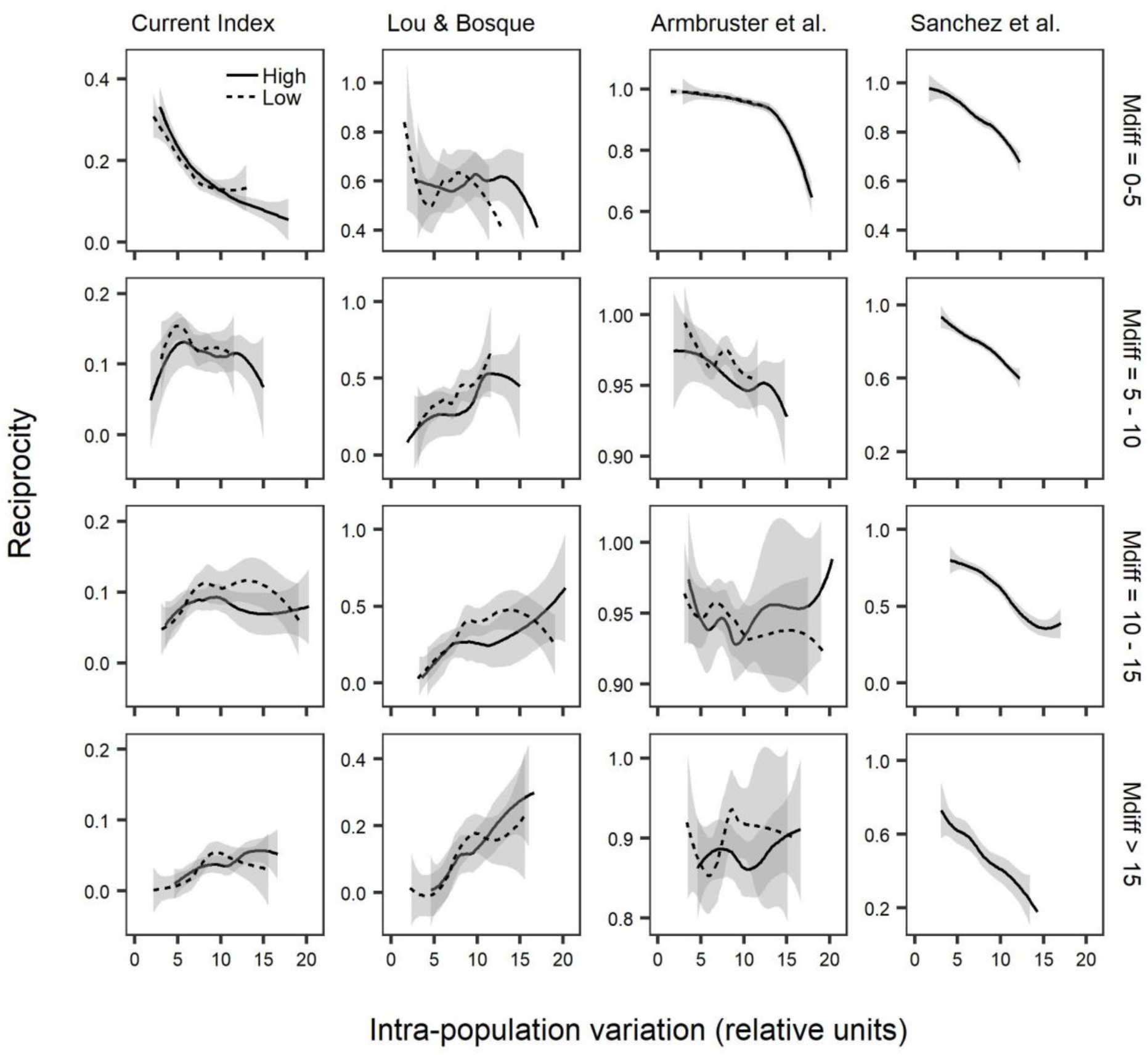
Comparison of the current index with previous indices over a range of intra-population variation in organ dimensions of naturally occurring heterostylous populations. The rows represent mean complementary sex organ heights (Mdiff) which are categorised as 0-5%, 5-10%, 10-15% and greater than 15%. Columns represents different indices. The solid and the dotted line represent high and low levels, respectively. The index proposed by Sanchez *et al*. (2013) is a composite value for the two levels in the flower. Values for Armbruster *et al*. (2017) are standardized and scaled to range between zero and one.

### Comparison of indices using naturally occurring populations

The indices proposed by (Richards and Koptur, 1993) modified to include intra-population variation, and the index proposed by Sanchez *et al*. (2013) always decreased with an increase in intra-population variation (Fig. 6). As seen in the simulations, the index proposed by Sanchez *et al*. (2013) showed high values of reciprocity even when difference in mean complementary sex organ heights is high (above 10%) and intra-population variation is negligible. This is contrary to expectations that under these circumstances there should be low overlap between the distribution of anther and stigma heights of the complementary morphs, and therefore low values for reciprocity. The estimated values of reciprocity from Lau and Bosque (2003) decreased with an increase in intra-population variation when the difference in mean complementary sex organ heights is low. When the difference in mean complementary sex organ heights is high, it increased with increasing intra-population variation. The index proposed by Armbruster *et al*. (2017) showed very little difference in reciprocity with an increase in intra-population variation in all the categories of difference in mean complementary sex organ heights.

### Do reciprocity indices predict reproductive success?

Data was extracted for fruit set from 52 populations (34 species), and for seed set 34 populations (27 species). Significant positive correlations were seen between index values and seed set for the current index, and the Armbruster et. al’s index in both the levels (Table 1). Modified Richards and Koptur’s index showed significant positive correlation for seed set in the higher, but not the low level. None of the indices showed a significant correlation with fruit set in both the levels.

**Table 1.**
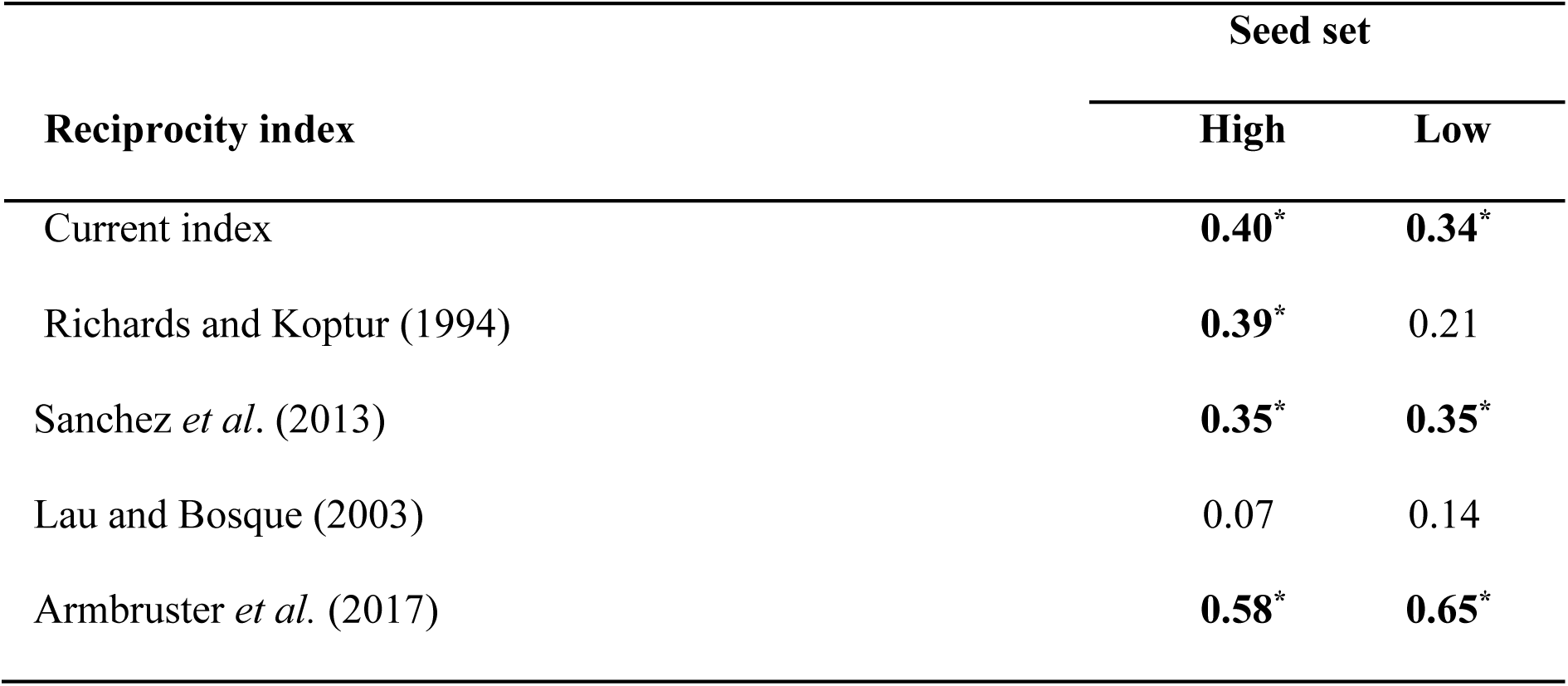
Spearman’s correlation coefficients for relationships between estimates of reciprocity and measures of reproductive success (seed set). High and Low refer to the seed set for the higher and lower sex-organ levels in the flower, respectively. Sample size: n = 34 populations. Significant correlations at *p* < 0.05 are denoted in bold and marked with *.

| Reciprocity index | Seed set |  |
| --- | --- | --- |
|  | High | Low |
| Current index | **0.40\*** | **0.34\*** |
| Richards and Koptur (1994) | **0.39\*** | 0.21 |
| Sanchez *et al.* (2013) | **0.35\*** | **0.35\*** |
| Lau and Bosque (2003) | 0.07 | 0.14 |
| Armbruster *et al.* (2017) | **0.58\*** | **0.65\*** |

## DISCUSSION

The results from the simulated populations demonstrated that when the difference in mean anther and stigma height of complementary morphs is zero or very small, increasing intra-population variation in sex organ heights resulted in a monotonous decrease in reciprocity as quantified using the index developed in this study. However, when the difference in mean complementary anther and stigma height is higher, reciprocity initially increased with increasing variation, reached a peak and then decreased. We also observed this when we examined the relationship between reciprocity and intrapopulation variation with data from naturally occurring heterostylous populations. These changes in reciprocity as a function of changing intra-population variation and difference in mean complementary sex organ heights are not captured by other commonly used reciprocity indices. These results are contrary to our current understanding that increasing variation in sex organ heights should always result in a decrease in reciprocity.

Reciprocity indices are widely used to understand the functional consequences of variation in sex organ position for legitimate pollen transfer between complementary morphs of heterostylous species (Pailler and Thompson, 1997; Faivre and McDade, 2001; Ferrero *et al*., 2009; Consolaro *et al*., 2011). These indices are utilized for understanding plant-pollinator interactions, pollen flow, fruit set, and the general reproductive biology of heterostylous plant populations (Brys *et al*., 2008; Ferrero, Castro, *et al*., 2011; Santos-Gally *et al*., 2013), and estimates of reciprocity indices have been applied to the management and conservation of species (Meeus *et al*., 2011; Casazza *et al*., 2013; Aronne *et al*., 2014; Chen *et al*., 2014). Reciprocity indices have contributed significantly to our understanding evolution of sexual systems and floral polymorphisms (Sosenski *et al*., 2010; Baena-Díaz *et al*., 2012; Yuan *et al*., 2017), and estimates of reciprocity between species have also been used to assess the potential for hybridization, and conversely the reproductive isolation between species (Zhu *et al*., 2009; Keller *et al*., 2012, 2016; Huang *et al*., 2015). Finally, quantitative estimates of sex organ positions and measures of reciprocity have been used to define the floral polymorphic status of populations (Dulberger, 1973; Richards and Koptur, 1993; Baker, 2000), and used as a taxonomic tool (Eiten, 1963; Selvi, 1998; Esteves and Vicentini, 2013).

We found that intra-population variation in sex organ heights and differences in mean heights between complementary sex organs can be substantial in heterostylous populations. Given this it is important to understand how combinations of intra-population variation and difference in mean complementary sex organ heights might alter reciprocity. The commonly used reciprocity indices, Richard and Koptur’s and Sanchez’s indices, and an index recently proposed by Armbruster failed to capture the loss of reciprocity that should result from increased differences in mean complementary sex organ heights, and the increase in reciprocity that is expected with increasing intrapopulation variation when there is substantial differences in mean complementary sex organ heights. The current index was able to account for these expected changes in reciprocity with changes in the distribution of anthers and stigmas. While with Lau and Bosque’s index we observed the expected increase in reciprocity with increasing intrapopulation variation, this index remained problematic because estimates of reciprocity continued to increase and ultimately saturate with increasing intrapopulation variation.

How mismatches ultimately translate to the amount of pollen transferred between complementary sex organs will depend on pick-up from the anthers, deposition on the pollinator body, redistribution during flight, and ultimately deposition onto the stigma, but empirical evidence for such patterns are very limited. Studies have shown that spatial distribution of pollen on the body, or proboscis of long-tongued pollinators (Levin and Berube, 1972; Courtney *et al*., 1982; Washitan *et al*., 1994; Harder and Wilson, 1998; Keller *et al*., 2014; Deschepper *et al*., 2018) differed for pollen from different morphs and therefore likely important for the reciprocal transfer to complementary sex organs. However, these and other studies also show substantial variability in the precision of pick up and deposition of pollen, and redistribution on the pollinator body during flight post pick-up, and this weakens the case for effective legitimate pollen transfer in heterostylous species. Given this lack of empirical evidence, we assumed a simple concave decreasing function for converting values of mismatch to a pollen transfer probability. We showed that relaxing this assumption, and testing other decreasing functions did not change the important qualitative nature of our results. However, there is an urgent need to better understand patterns of pollen pick-up and deposition, and spatial distribution on the pollinator.

Congruent with the results we observed in the simulated populations, we observed that the different indices yielded very different estimates of reciprocity for naturally occurring populations. With the exception of (Lau and Bosque, 2003), the other indices consistently overestimated reciprocity in natural plant populations in comparison to the current index. This is likely because the other indices were not sensitive to decreases in reciprocity that should result from increased differences in mean complementary sex organ heights. As such the current index was more sensitive and better at discriminating between reciprocity estimates for species.

The current index and the indices proposed by Sanchez *et al*. (2013) and Armbruster *et al*. (2017) showed significant positive correlation with seed set. Reciprocity indices are used to understand the efficiency of legitimate pollen transfer and consequently reproductive success in heterostylous plant populations. Our results show that reciprocity indices can predict reproductive success of naturally occurring heterostylous plant populations. However, it is also important to recognize that fruit set and seed set are influenced by a number of other important factors like resource limitation, resource allocation, self-incompatibility etc. (Charlesworth, 1989).

The reciprocity index proposed here exhibited markedly different quantitative and qualitative behaviour from previous indices as a response to increasing intra-population variation in sex organ heights and difference in mean complementary sex organ heights. Challenging current understanding, our results suggest that increasing intra-population variation in sex organ heights can result in an increase in reciprocity in heterostylous populations. This might explain how heterostylous systems exhibit, and tolerate high amounts of intra-population variation in sex organ heights. Such variation can facilitate the stabilisation and perpetuation of imperfectly reciprocal states that are in the process of evolving towards perfect heterostyly.

## Acknowledgements

We thank Sudipta Tung for help with the simulations. S.G. was supported by a Fellowship from Indian Institute of Science Education and Research, Pune.

## Supplementary information

**Figure S1:**
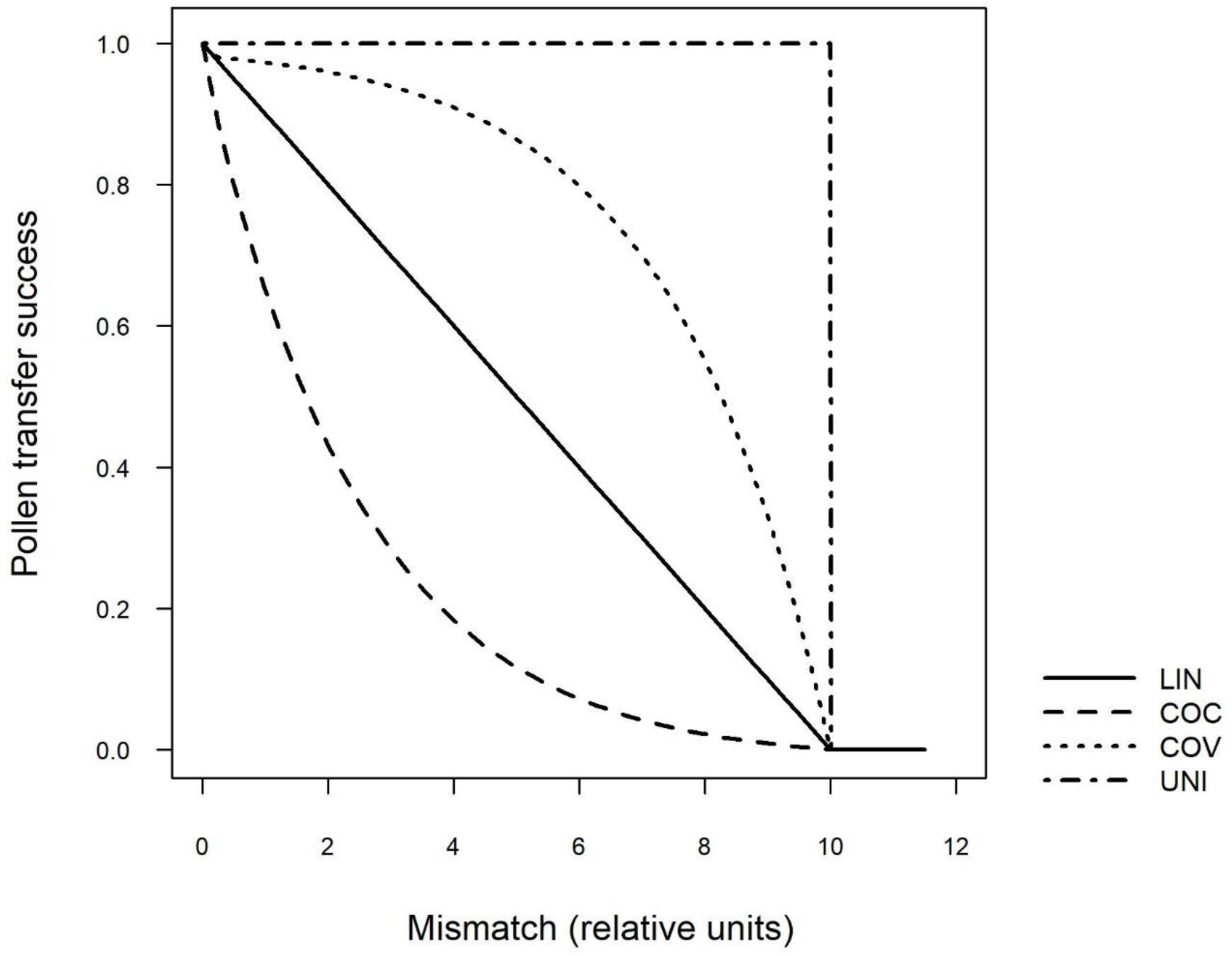
The different pollen transfer functions examined: COC - concave decreasing; LIN - linear decreasing; COV - Convex decreasing; and, UNI - Uniform step function. This function was used to convert mismatch between complementary organ heights to pollen transfer success estimates. We assumed a critical value of mismatch beyond which there was no pollen transfer. The formula used for the convex, linear and concave functions are: 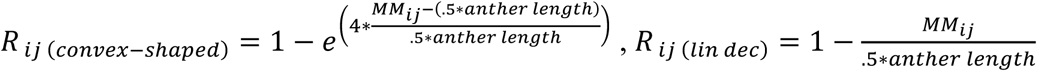 and 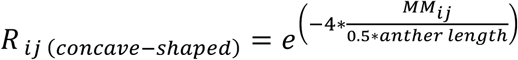 respectively. Here, *R_ij_* is pollen transfer success, and M*_ij_* the mismatch between the *i*th anther and *j*th stigma of complementary morphs.

**Figure S2:**
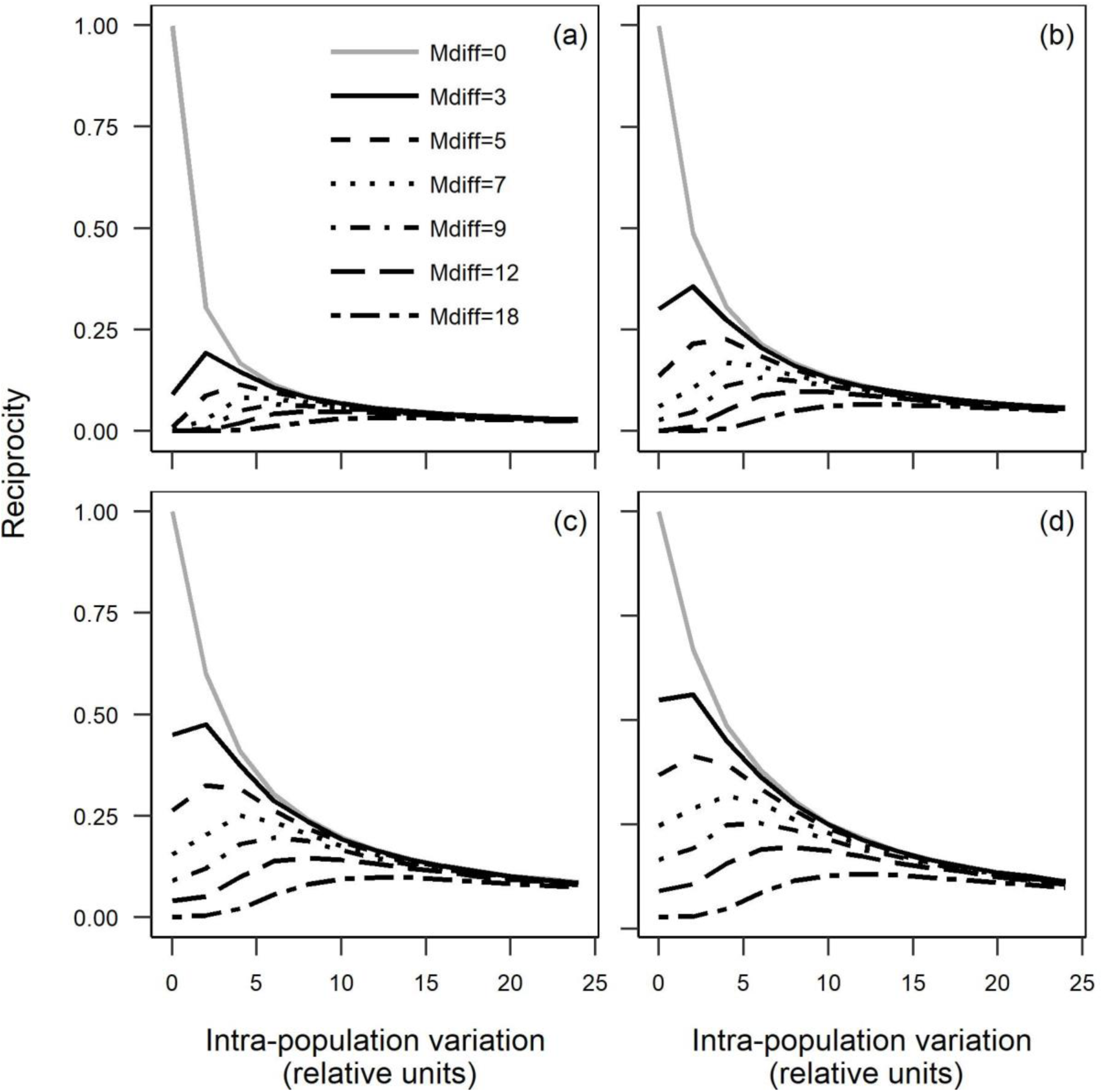
Comparison of reciprocity estimates when the the critical value of mismatch at which pollen transfer success becomes zero is set at: (a) 5%, (b) 10%, (c) 15% and (d) 20% of the mean height of the high level sex organs using the concave shaped function. Reciprocity estimates for the simulated populations as a function of increasing intra-population variation in anther and stigma heights, for a range of values for difference in mean sex organ heights (Mdiff; shown by the different curves).

**Table S1:** Comparison of the proposed index (COC-concave shaped pollen transfer success fucntion) with modified Richards and Koptur - R & K (Sanchez *et al.*, 2008), Sanchez - SAN (Sanchez *et al*., 2013), Lou and Bosque – L & B (Lou and Bosque, 2003) & Armbruster – ARM (Armbruster *et al*., 2017) using extracted individual level empirical data. Sanchez’s index represents a composite value for both the levels of organ positions and is compared to an average value of index of both the high and low levels for all other indices. Fruit-set and seed-set are represented as percentage per flower and per ovule respectively. Reference denotes the serial number for the study from which data was extracted as per the bibliography in the supplementary references. The species names presented here are accepted names according to The Plant List (2013) (Version 1.1. Published on the Internet; http://www.theplantlist.org/ (accessed 17 April 2019)

| Species | Level | Reciprocity index values |  |  |  |  | Reproductive success |  | Reference |
| --- | --- | --- | --- | --- | --- | --- | --- | --- | --- |
|  |  | COC | R & K | SAN | L & B | ARM | Fruit set | Seed set |  |
| *Arcytophyllum aristatum* | High | 0.22 | 0.96 | - | 1.00 | 0.00 | - | 53.80 | 1 |
|  | Low | 0.15 | 0.94 | - | 0.33 | 0.00 | - | 79.10 |  |
|  | Average | 0.19 | 0.95 | 0.91 | 0.67 | 0.00 | - | - |  |
| *Arcytophyllum capitatum* | High | 0.10 | 0.91 | - | 0.56 | 0.03 | - | 43.80 | 1 |
|  | Low | 0.18 | 0.92 | - | 0.82 | 0.01 | - | 50.00 |  |
|  | Average | 0.14 | 0.91 | 0.59 | 0.69 | 0.02 | - | - |  |
| *Arcytophyllum ciliolatum* | High | 0.12 | 0.92 | - | 0.75 | 0.01 | - | 58.00 | 1 |
|  | Low | 0.16 | 0.90 | - | 0.92 | 0.00 | - | 65.10 |  |
|  | Average | 0.14 | 0.91 | 0.72 | 0.83 | 0.01 | - | - |  |
| *Arcytophyllum filiforme* | High | 0.07 | 0.87 | - | 0.70 | 0.02 | - | 50.50 | 1 |
|  | Low | 0.13 | 0.87 | - | 0.60 | 0.01 | - | 69.30 |  |
|  | Average | 0.10 | 0.87 | 0.41 | 0.65 | 0.02 | - | - |  |
| *Arcytophyllum lavarum* | High | 0.23 | 0.94 | - | 0.82 | 0.01 | - | 88.00 | 1 |
|  | Low | 0.18 | 0.86 | - | 0.60 | 0.01 | - | 90.10 |  |
|  | Average | 0.21 | 0.90 | 0.67 | 0.71 | 0.01 | - | - |  |
| *Arcytophyllum macbridei* | High | 0.12 | 0.92 | - | 0.64 | 0.03 | - | 50.20 | 1 |
|  | Low | 0.19 | 0.93 | - | 0.53 | 0.01 | - | 65.00 |  |
|  | Average | 0.15 | 0.93 | 0.76 | 0.58 | 0.02 | - | - |  |

Table S1 continued.
| Species | Level | Reciprocity index values |  |  |  |  | Reproductive success |  | Reference |
| --- | --- | --- | --- | --- | --- | --- | --- | --- | --- |
|  |  | COC | R & K | SAN | L & B | ARM | Fruit set | Seed set |  |
| *Arcytophyllum macbridei* | High | 0.04 | 0.91 | - | 0.40 | 0.03 | - | 54.10 | 1 |
|  | Low | 0.26 | 0.93 | - | 0.87 | 0.01 | - | 54.20 |  |
|  | Average | 0.15 | 0.92 | 0.75 | 0.63 | 0.02 | - | - |  |
| *Arcytophyllum rivetii* | High | 0.15 | 0.93 | - | 0.90 | 0.01 | - | 85.70 | 1 |
|  | Low | 0.19 | 0.91 | - | 0.73 | 0.01 | - | 89.30 |  |
|  | Average | 0.17 | 0.92 | 0.70 | 0.81 | 0.01 | - | - |  |
| *Arcytophyllum setosum* | High | 0.09 | 0.93 | - | 0.60 | 0.02 | - | 82.80 | 1 |
|  | Low | 0.10 | 0.86 | - | 0.40 | 0.02 | - | 84.50 |  |
|  | Average | 0.09 | 0.89 | 0.66 | 0.50 | 0.02 | - | - |  |
| *Arcytophyllum thymifolium* | High | 0.11 | 0.92 | - | 0.57 | 0.01 | - | 82.30 | 1 |
|  | Low | 0.15 | 0.91 | - | 0.71 | 0.01 | - | 97.70 |  |
|  | Average | 0.13 | 0.91 | 0.70 | 0.64 | 0.01 | - | - |  |
| *Arcytophyllum vernicosum* | High | 0.12 | 0.92 | - | 0.71 | 0.01 | - | 62.60 | 1 |
|  | Low | 0.15 | 0.91 | - | 0.58 | 0.01 | - | 85.80 |  |
|  | Average | 0.14 | 0.91 | 0.69 | 0.65 | 0.01 | - | - |  |
| *Bouvardia ternifolia* | High | 0.14 | 0.95 | - | 0.12 | 15.33 | - | - | 2 |
|  | Low | 0.16 | 0.94 | - | 0.12 | 10.90 | - | - |  |
|  | Average | 0.15 | 0.94 | 0.86 | 0.12 | 13.11 | - | - |  |
| *Carapichea ipecacuanha* | High | 0.11 | 0.93 | - | 0.55 | 1.10 | 25.00 | - | 3 |
|  | Low | 0.19 | 0.91 | - | 0.28 | 0.47 | 16.70 | - |  |
|  | Average | 0.15 | 0.92 | 0.79 | 0.42 | 0.79 | - | - |  |

Table S1 continued.
| Species | Level | Reciprocity index values |  |  |  |  | Reproductive success |  | Reference |
| --- | --- | --- | --- | --- | --- | --- | --- | --- | --- |
|  |  | COC | R & K | SAN | L & B | ARM | Fruit set | Seed set |  |
| *Carapichea ipecacuanha* | High | 0.18 | 0.95 | - | 0.48 | 0.96 | - | - | 3 |
|  | Low | 0.15 | 0.91 | - | 0.42 | 0.62 | - | - |  |
|  | Average | 0.16 | 0.93 | 0.82 | 0.45 | 0.79 | - | - |  |
| *Carapichea ipecacuanha* | High | 0.12 | 0.92 | - | 0.61 | 1.57 | - | - | 3 |
|  | Low | 0.17 | 0.90 | - | 0.52 | 0.69 | - | - |  |
|  | Average | 0.15 | 0.91 | 0.72 | 0.56 | 1.13 | - | - |  |
| *Chassalia corallioides* | High | 0.09 | 0.93 | - | 0.22 | 7.55 | 58.60 | - | 4 |
|  | Low | 0.01 | 0.77 | - | 0.05 | 29.83 | 0.00 | - |  |
|  | Average | 0.05 | 0.85 | 0.46 | 0.13 | 18.69 | - | - |  |
| *Chassalia corallioides* | High | 0.07 | 0.90 | - | 0.14 | 15.12 | 45.90 | - | 4 |
|  | Low | 0.01 | 0.68 | - | 0.00 | 42.63 | 0.00 | - |  |
|  | Average | 0.04 | 0.79 | -0.02 | 0.07 | 28.87 | - | - |  |
| *Chassalia corallioides* | High | 0.10 | 0.94 | - | 0.33 | 4.09 | 36.60 | - | 4 |
|  | Low | 0.04 | 0.82 | - | 0.08 | 14.02 | 0.00 | - |  |
|  | Average | 0.07 | 0.88 | 0.57 | 0.21 | 9.05 | - | - |  |
| *Chassalia corallioides* | High | 0.11 | 0.94 | - | 0.32 | 5.24 | 28.30 | - | 4 |
|  | Low | 0.02 | 0.81 | - | 0.04 | 19.38 | 0.00 | - |  |
|  | Average | 0.07 | 0.87 | 0.53 | 0.18 | 12.31 | - | - |  |
| *Chassalia corallioides* | High | 0.09 | 0.94 | - | 0.20 | 8.45 | 29.90 | - | 4 |
|  | Low | 0.01 | 0.82 | - | 0.03 | 21.77 | 0.00 | - |  |
|  | Average | 0.05 | 0.88 | 0.64 | 0.12 | 15.11 | - | - |  |

Table S1 continued.
| Species | Level | Reciprocity index values |  |  |  |  | Reproductive success |  | Reference |
| --- | --- | --- | --- | --- | --- | --- | --- | --- | --- |
|  |  | COC | R & K | SAN | L & B | ARM | Fruit set | Seed set |  |
| *Cordia macrocephala* | High | 0.09 | 0.93 | - | 0.14 | 4.84 | 32.00 | - | 5 |
|  | Low | 0.06 | 0.89 | - | 0.18 | 6.67 | 30.00 | - |  |
|  | Average | 0.08 | 0.91 | 0.73 | 0.16 | 5.75 | - | - |  |
| *Coussarea croceoides* | High | 0.21 | 0.93 | - | 0.43 | 6.67 | - | - | 6 |
|  | Low | 0.18 | 0.88 | - | 0.43 | 9.13 | - | - |  |
|  | Average | 0.19 | 0.90 | 0.67 | 0.43 | 7.90 | - | - |  |
| *Coussarea platyphylla* | High | 0.04 | 0.89 | - | 0.04 | 130.44 | 85.00 | - | 6 |
|  | Low | 0.03 | 0.83 | - | 0.04 | 164.84 | 90.00 | - |  |
|  | Average | 0.03 | 0.86 | 0.37 | 0.04 | 147.64 | - | - |  |
| *Damnacanthus macrophyllus* | High | 0.08 | 0.93 | - | 0.18 | 4.21 | - | - | 7 |
|  | Low | 0.09 | 0.90 | - | 0.27 | 3.53 | - | - |  |
|  | Average | 0.09 | 0.92 | 0.75 | 0.23 | 3.87 | - | - |  |
| *Damnacanthus major* | High | 0.06 | 0.90 | - | 0.10 | 9.71 | - | - | 7 |
|  | Low | 0.09 | 0.90 | - | 0.17 | 4.23 | - | - |  |
|  | Average | 0.07 | 0.90 | 0.59 | 0.13 | 6.97 | - | - |  |
| *Declieuxia fruticosa* | High | 0.10 | 0.92 | - | 0.49 | 1.25 | 86.70 | - | 8 |
|  | Low | 0.05 | 0.85 | - | 0.23 | 1.44 | 53.30 | - |  |
|  | Average | 0.08 | 0.89 | 0.61 | 0.36 | 1.35 | - | - |  |
| *Faramea occidentalis* | High | 0.18 | 0.94 | - | 0.80 | 0.04 | 28.00 | - | 9 |
|  | Low | 0.07 | 0.83 | - | 0.31 | 0.10 | 20.00 | - |  |
|  | Average | 0.13 | 0.88 | 0.61 | 0.56 | 0.07 | - | - |  |

Table S1 continued.
| Species | Level | Reciprocity index values |  |  |  |  | Reproductive success |  | Reference |
| --- | --- | --- | --- | --- | --- | --- | --- | --- | --- |
|  |  | COC | R & K | SAN | L & B | ARM | Fruit set | Seed set |  |
| *Faramea suerrensis* | High | 0.10 | 0.92 | - | 0.19 | 2.10 | - | - | 10 |
|  | Low | 0.00 | 0.72 | - | 0.00 | 5.71 | - | - |  |
|  | Average | 0.05 | 0.82 | 0.51 | 0.10 | 3.91 | - | - |  |
| *Gaertnera vaginata* | High | 0.06 | 0.92 | - | 0.17 | 17.79 | - | - | 11 |
|  | Low | 0.06 | 0.90 | - | 0.08 | 12.40 | - | - |  |
|  | Average | 0.06 | 0.91 | 0.75 | 0.12 | 15.10 | - | - |  |
| *Gaertnera vaginata* | High | 0.06 | 0.92 | - | 0.17 | 18.13 | - | - | 11 |
|  | Low | 0.10 | 0.92 | - | 0.25 | 9.33 | - | - |  |
|  | Average | 0.08 | 0.92 | 0.75 | 0.21 | 13.73 | - | - |  |
| *Gaertnera vaginata* | High | 0.07 | 0.91 | - | 0.09 | 22.60 | - | - | 11 |
|  | Low | 0.13 | 0.91 | - | 0.09 | 9.00 | - | - |  |
|  | Average | 0.10 | 0.91 | 0.69 | 0.09 | 15.80 | - | - |  |
| *Gaertnera vaginata* | High | 0.03 | 0.89 | - | 0.08 | 29.48 | - | - | 11 |
|  | Low | 0.16 | 0.93 | - | 0.33 | 5.92 | - | - |  |
|  | Average | 0.10 | 0.91 | 0.68 | 0.21 | 17.70 | - | - |  |
| *Gaertnera vaginata* | High | 0.05 | 0.92 | - | 0.04 | 17.83 | - | - | 11 |
|  | Low | 0.09 | 0.89 | - | 0.17 | 15.21 | - | - |  |
|  | Average | 0.07 | 0.91 | 0.75 | 0.11 | 16.52 | - | - |  |
| *Gaertnera vaginata* | High | 0.06 | 0.93 | - | 0.13 | 14.12 | - | - | 11 |
|  | Low | 0.16 | 0.94 | - | 0.29 | 5.17 | - | - |  |
|  | Average | 0.11 | 0.94 | 0.86 | 0.21 | 9.65 | - | - |  |

Table S1 continued.
| Species | Level | Reciprocity index values |  |  |  |  | Reproductive success |  | Reference |
| --- | --- | --- | --- | --- | --- | --- | --- | --- | --- |
|  |  | COC | R & K | SAN | L & B | ARM | Fruit set | Seed set |  |
| *Gaertnera vaginata* | High | 0.10 | 0.93 | - | 0.16 | 14.45 | 29.00 | - | 11 |
|  | Low | 0.06 | 0.89 | - | 0.04 | 11.64 | 21.00 | - |  |
|  | Average | 0.08 | 0.91 | 0.76 | 0.10 | 13.04 | - | - |  |
| *Gaertnera vaginata* | High | 0.07 | 0.91 | - | 0.17 | 18.52 | 49.00 | - | 11 |
|  | Low | 0.13 | 0.91 | - | 0.28 | 7.27 | 39.00 | - |  |
|  | Average | 0.10 | 0.91 | 0.71 | 0.23 | 12.90 | - | - |  |
| *Glandora diffusa* | High | 0.07 | 0.92 | - | 0.70 | 4.55 | - | - | 12 |
|  | Low | 0.06 | 0.88 | - | 0.63 | 3.91 | - | - |  |
|  | Average | 0.07 | 0.90 | 0.66 | 0.67 | 4.23 | - | - |  |
| *Glandora diffusa* | High | 0.09 | 0.94 | - | 0.88 | 1.90 | - | - | 12 |
|  | Low | 0.12 | 0.93 | - | 0.94 | 0.98 | - | - |  |
|  | Average | 0.10 | 0.93 | 0.82 | 0.91 | 1.44 | - | - |  |
| *Glandora moroccana* | High | 0.08 | 0.93 | - | 0.74 | 5.48 | 47.00 | 12.00 | 12, 13 |
|  | Low | 0.12 | 0.92 | - | 0.80 | 2.72 | 76.00 | 19.00 |  |
|  | Average | 0.10 | 0.92 | 0.74 | 0.77 | 4.10 | - | - |  |
| *Glandora moroccana* | High | 0.09 | 0.93 | - | 0.69 | 5.51 | - | - | 12 |
|  | Low | 0.13 | 0.92 | - | 0.76 | 2.04 | - | - |  |
|  | Average | 0.11 | 0.93 | 0.77 | 0.72 | 3.78 | - | - |  |
| *Glandora oleifolia* | High | 0.04 | 0.92 | - | 0.17 | 6.31 | - | - | 12 |
|  | Low | 0.13 | 0.90 | - | 0.90 | 2.58 | - | - |  |
|  | Average | 0.08 | 0.91 | 0.72 | 0.53 | 4.44 | - | - |  |

Table S1 continued.
| Species | Level | Reciprocity index values |  |  |  |  | Reproductive success |  | Reference |
| --- | --- | --- | --- | --- | --- | --- | --- | --- | --- |
|  |  | COC | R & K | SAN | L & B | ARM | Fruit set | Seed set |  |
| *Glandora oleifolia* | High | 0.09 | 0.94 | - | 0.73 | 3.45 | 27.00 | 7.00 | 12, 13 |
|  | Low | 0.08 | 0.90 | - | 0.31 | 3.01 | 30.00 | 8.00 |  |
|  | Average | 0.09 | 0.92 | 0.81 | 0.52 | 3.23 | - | - |  |
| *Glandora rosmarinifolia* | High | 0.10 | 0.95 | - | 0.58 | 1.88 | - | - | 12 |
|  | Low | 0.08 | 0.92 | - | 0.36 | 1.44 | - | - |  |
|  | Average | 0.09 | 0.93 | 0.85 | 0.47 | 1.66 | - | - |  |
| *Glandora rosmarinifolia* | High | 0.06 | 0.93 | - | 0.48 | 5.95 | - | - | 12 |
|  | Low | 0.10 | 0.92 | - | 0.63 | 2.32 | - | - |  |
|  | Average | 0.08 | 0.92 | 0.75 | 0.55 | 4.14 | - | - |  |
| *Goniolimon italicum* | High | 0.08 | 0.94 | - | 0.07 | 0.95 | - | - | 14 |
|  | Low | 0.21 | 0.96 | - | 0.36 | 0.44 | - | - |  |
|  | Average | 0.15 | 0.95 | 0.91 | 0.22 | 0.69 | - | - |  |
| *Houstonia caerulea* | High | 0.04 | 0.88 | - | 0.28 | 2.63 | - | - | 15 |
|  | Low | 0.10 | 0.86 | - | 0.28 | 0.83 | - | - |  |
|  | Average | 0.07 | 0.87 | 0.47 | 0.28 | 1.73 | - | - |  |
| *Houstonia caerulea* | High | 0.05 | 0.87 | - | 0.28 | 3.33 | - | - | 15 |
|  | Low | 0.22 | 0.92 | - | 0.59 | 0.31 | - | - |  |
|  | Average | 0.13 | 0.89 | 0.38 | 0.43 | 1.82 | - | - |  |
| *Houstonia caerulea* | High | 0.06 | 0.89 | - | 0.31 | 3.02 | - | - | 15 |
|  | Low | 0.14 | 0.89 | - | 0.39 | 0.77 | - | - |  |
|  | Average | 0.10 | 0.89 | 0.51 | 0.35 | 1.89 | - | - |  |

Table S1 continued.
| Species | Level | Reciprocity index values |  |  |  |  | Reproductive success |  | Reference |
| --- | --- | --- | --- | --- | --- | --- | --- | --- | --- |
|  |  | COC | R & K | SAN | L & B | ARM | Fruit set | Seed set |  |
| *Houstonia longifolia* | High | 0.06 | 0.91 | - | 0.26 | 1.14 | - | - | 15 |
|  | Low | 0.23 | 0.93 | - | 0.71 | 0.14 | - | - |  |
|  | Average | 0.15 | 0.92 | 0.72 | 0.48 | 0.64 | - | - |  |
| *Houstonia longifolia* | High | 0.05 | 0.91 | - | 0.18 | 1.10 | - | - | 15 |
|  | Low | 0.31 | 0.95 | - | 0.52 | 0.09 | - | - |  |
|  | Average | 0.18 | 0.93 | 0.75 | 0.35 | 0.60 | - | - |  |
| *Houstonia procumbens* | High | 0.13 | 0.94 | - | 0.56 | 1.84 | - | - | 15 |
|  | Low | 0.16 | 0.92 | - | 0.50 | 1.13 | - | - |  |
|  | Average | 0.14 | 0.93 | 0.81 | 0.53 | 1.49 | - | - |  |
| *Houstonia procumbens* | High | 0.11 | 0.93 | - | 0.75 | 2.90 | - | - | 15 |
|  | Low | 0.13 | 0.91 | - | 0.32 | 2.44 | - | - |  |
|  | Average | 0.12 | 0.92 | 0.71 | 0.54 | 2.67 | - | - |  |
| *Houstonia procumbens* | High | 0.15 | 0.95 | - | 0.30 | 1.82 | - | - | 15 |
|  | Low | 0.07 | 0.89 | - | 0.26 | 2.64 | - | - |  |
|  | Average | 0.11 | 0.92 | 0.79 | 0.28 | 2.23 | - | - |  |
| *Jasminum fruticans* | High | 0.01 | 0.84 | - | 0.00 | 16.20 | - | 9.26 | 16, 17 |
|  | Low | 0.09 | 0.90 | - | 0.22 | 2.70 | - | 9.41 |  |
|  | Average | 0.05 | 0.87 | 0.45 | 0.11 | 9.45 | - | - |  |
| *Jasminum fruticans* | High | 0.04 | 0.89 | - | 0.09 | 8.74 | - | 8.56 | 16, 17 |
|  | Low | 0.04 | 0.84 | - | 0.13 | 6.71 | - | 10.37 |  |
|  | Average | 0.04 | 0.86 | 0.43 | 0.11 | 7.73 | - | - |  |

Table S1 continued.
| Species | Level | Reciprocity index values |  |  |  |  | Reproductive success |  | Reference |
| --- | --- | --- | --- | --- | --- | --- | --- | --- | --- |
|  |  | COC | R & K | SAN | L & B | ARM | Fruit set | Seed set |  |
| *Jasminum fruticans* | High | 0.00 | 0.84 | - | 0.00 | 12.32 | - | - | 16 |
|  | Low | 0.08 | 0.90 | - | 0.19 | 2.32 | - | - |  |
|  | Average | 0.04 | 0.87 | 0.62 | 0.09 | 7.32 | - | - |  |
| *Jasminum fruticans* | High | 0.04 | 0.87 | - | 0.11 | 10.74 | - | 10.48 | 16, 17 |
|  | Low | 0.10 | 0.88 | - | 0.32 | 5.11 | - | 10.64 |  |
|  | Average | 0.07 | 0.88 | 0.38 | 0.22 | 7.92 | - | - |  |
| *Jasminum fruticans* | High | 0.00 | 0.84 | - | 0.00 | 12.01 | - | - | 16 |
|  | Low | 0.21 | 0.95 | - | 0.36 | 0.60 | - | - |  |
|  | Average | 0.11 | 0.90 | 0.60 | 0.18 | 6.30 | - | - |  |
| *Jasminum fruticans* | High | 0.00 | 0.84 | - | 0.00 | 14.55 | - | 8.94 | 16, 17 |
|  | Low | 0.14 | 0.92 | - | 0.25 | 2.15 | - | 17.23 |  |
|  | Average | 0.07 | 0.88 | 0.50 | 0.13 | 8.35 | - | - |  |
| *Jasminum fruticans* | High | 0.02 | 0.86 | - | 0.09 | 11.35 | - | - | 16 |
|  | Low | 0.14 | 0.91 | - | 0.45 | 1.79 | - | - |  |
|  | Average | 0.08 | 0.89 | 0.47 | 0.27 | 6.57 | - | - |  |
| *Jasminum fruticans* | High | 0.02 | 0.84 | - | 0.08 | 14.06 | - | 15.43 | 16, 17 |
|  | Low | 0.16 | 0.92 | - | 0.50 | 1.54 | - | 11.97 |  |
|  | Average | 0.09 | 0.88 | 0.39 | 0.29 | 7.80 | - | - |  |
| *Jasminum fruticans* | High | 0.15 | 0.94 | - | 0.40 | 3.36 | - | - | 16 |
|  | Low | 0.13 | 0.92 | - | 0.22 | 2.16 | - | - |  |
|  | Average | 0.14 | 0.93 | 0.80 | 0.31 | 2.76 | - | - |  |

Table S1 continued.
| Species | Level | Reciprocity index values |  |  |  |  | Reproductive success |  | Reference |
| --- | --- | --- | --- | --- | --- | --- | --- | --- | --- |
|  |  | COC | R & K | SAN | L & B | ARM | Fruit set | Seed set |  |
| *Jasminum fruticans* | High | 0.06 | 0.90 | - | 0.14 | 6.68 | - | - | 16 |
|  | Low | 0.10 | 0.89 | - | 0.17 | 2.69 | - | - |  |
|  | Average | 0.08 | 0.89 | 0.60 | 0.15 | 4.68 | - | - |  |
| *Jasminum fruticans* | High | 0.12 | 0.94 | - | 0.38 | 3.05 | - | - | 16 |
|  | Low | 0.08 | 0.86 | - | 0.29 | 5.55 | - | - |  |
|  | Average | 0.10 | 0.90 | 0.66 | 0.34 | 4.30 | - | - |  |
| *Kalmiopsis fragrans* | High | 0.04 | 0.92 | - | 0.10 | 6.80 | 64.29 | - | 18 |
|  | Low | 0.02 | 0.81 | - | 0.05 | 12.17 | 64.29 | - |  |
|  | Average | 0.03 | 0.86 | 0.52 | 0.07 | 9.48 | - | - |  |
| *Linum aretioides* | High | 0.00 | 0.82 | - | 0.00 | 5.10 | 71.06 | - | 19 |
|  | Low | 0.12 | 0.90 | - | 0.13 | 0.56 | 73.02 | - |  |
|  | Average | 0.06 | 0.86 | 0.35 | 0.07 | 2.83 | - | - |  |
| *Linum aretioides* | High | 0.06 | 0.84 | - | 0.18 | 5.73 | - | - | 19 |
|  | Low | 0.06 | 0.83 | - | 0.23 | 2.92 | - | - |  |
|  | Average | 0.06 | 0.84 | -0.09 | 0.20 | 4.33 | - | - |  |
| *Linum campanulatum* | High | 0.17 | 0.95 | - | 0.48 | 2.48 | - | - | 16 |
|  | Low | 0.09 | 0.89 | - | 0.13 | 5.67 | - | - |  |
|  | Average | 0.13 | 0.92 | 0.80 | 0.30 | 4.07 | - | - |  |
| *Linum suffruticosum* | High | 0.15 | 0.93 | - | 0.34 | 1.52 | - | - | 20 |
|  | Low | 0.13 | 0.88 | - | 0.44 | 1.39 | - | - |  |
|  | Average | 0.14 | 0.90 | 0.65 | 0.39 | 1.46 | - | - |  |

Table S1 continued.
| Species | Level | Reciprocity index values |  |  |  |  | Reproductive success |  | Reference |
| --- | --- | --- | --- | --- | --- | --- | --- | --- | --- |
|  |  | COC | R & K | SAN | L & B | ARM | Fruit set | Seed set |  |
| *Linum tenue* | High | 0.18 | 0.94 | - | 0.41 | 0.84 | - | - | 20 |
|  | Low | 0.00 | 0.72 | - | 0.00 | 3.72 | - | - |  |
|  | Average | 0.09 | 0.83 | 0.60 | 0.21 | 2.28 | - | - |  |
| *Linum tenuifolium* | High | 0.12 | 0.94 | - | 0.51 | 1.49 | - | - | 21 |
|  | Low | 0.13 | 0.93 | - | 0.64 | 1.16 | - | - |  |
|  | Average | 0.13 | 0.93 | 0.81 | 0.58 | 1.33 | - | - |  |
| *Luculia pinceana* | High | 0.19 | 0.96 | - | 0.27 | 7.51 | - | - | 22 |
|  | Low | 0.27 | 0.97 | - | 0.37 | 3.14 | - | - |  |
|  | Average | 0.23 | 0.96 | 0.94 | 0.32 | 5.33 | - | - |  |
| *Luculia pinceana* | High | 0.25 | 0.97 | - | 0.28 | 4.63 | - | - | 22 |
|  | Low | 0.24 | 0.96 | - | 0.20 | 4.14 | - | - |  |
|  | Average | 0.25 | 0.97 | 0.96 | 0.24 | 4.39 | - | - |  |
| *Luculia pinceana* | High | 0.39 | 0.98 | - | 0.40 | 1.57 | - | - | 22 |
|  | Low | 0.34 | 0.97 | - | 0.43 | 2.03 | - | - |  |
|  | Average | 0.36 | 0.98 | 0.98 | 0.42 | 1.80 | - | - |  |
| *Melochia nudiflora* | High | 0.08 | 0.93 | - | 0.30 | 1.99 | - | - | 23 |
|  | Low | 0.20 | 0.94 | - | 0.45 | 0.56 | - | - |  |
|  | Average | 0.14 | 0.94 | 0.85 | 0.38 | 1.27 | - | - |  |
| *Melochia pyramidata* | High | 0.14 | 0.94 | - | 0.91 | 0.47 | - | 40.05 | 23, 24 |
|  | Low | 0.10 | 0.90 | - | 1.00 | 0.83 | - | 30.97 |  |
|  | Average | 0.12 | 0.92 | 0.76 | 0.96 | 0.65 | - | - |  |

Table S1 continued.
| Species | Level | Reciprocity index values |  |  |  |  | Reproductive success |  | Reference |
| --- | --- | --- | --- | --- | --- | --- | --- | --- | --- |
|  |  | COC | R & K | SAN | L & B | ARM | Fruit set | Seed set |  |
| *Melochia savannarum* | High | 0.08 | 0.91 | - | 0.77 | 0.65 | - | - | 23 |
|  | Low | 0.02 | 0.80 | - | 0.39 | 0.97 | - | - |  |
|  | Average | 0.05 | 0.85 | 0.41 | 0.58 | 0.81 | - | - |  |
| *Melochia savannarum* | High | 0.06 | 0.88 | - | 0.80 | 1.79 | - | 15.66 | 23, 24 |
|  | Low | 0.08 | 0.87 | - | 0.53 | 1.00 | - | 7.13 |  |
|  | Average | 0.07 | 0.87 | 0.37 | 0.67 | 1.40 | - | - |  |
| *Melochia savannarum* | High | 0.11 | 0.92 | - | 0.91 | 2.80 | - | - | 23 |
|  | Low | 0.13 | 0.89 | - | 0.89 | 1.80 | - | - |  |
|  | Average | 0.12 | 0.91 | 0.69 | 0.90 | 2.30 | - | - |  |
| *Melochia tomentosa* | High | 0.08 | 0.92 | - | 0.25 | 3.53 | 80.00 | - | 25 |
|  | Low | 0.05 | 0.78 | - | 0.17 | 4.44 | 83.00 | - |  |
|  | Average | 0.06 | 0.85 | 0.49 | 0.21 | 3.99 | - | - |  |
| *Melochia tomentosa* | High | 0.07 | 0.88 | - | 0.62 | 3.90 | - | 9.59 | 23, 24 |
|  | Low | 0.15 | 0.92 | - | 1.00 | 0.86 | - | 22.96 |  |
|  | Average | 0.11 | 0.90 | 0.53 | 0.81 | 2.38 | - | - |  |
| *Melochia tomentosa* | High | 0.07 | 0.92 | - | 0.00 | 3.89 | 82.00 | - | 26 |
|  | Low | 0.28 | 0.95 | - | 0.30 | 0.66 | 78.00 | - |  |
|  | Average | 0.17 | 0.93 | 0.80 | 0.15 | 2.28 | - | - |  |
| *Melochia villosa* | High | 0.08 | 0.92 | - | 0.81 | 1.70 | - | 6.83 | 23, 24 |
|  | Low | 0.08 | 0.90 | - | 0.38 | 1.22 | - | 16.48 |  |
|  | Average | 0.08 | 0.91 | 0.73 | 0.59 | 1.46 | - | - |  |

Table S1 continued.
| Species | Level | Reciprocity index values |  |  |  |  | Reproductive success |  | Reference |
| --- | --- | --- | --- | --- | --- | --- | --- | --- | --- |
|  |  | COC | R & K | SAN | L & B | ARM | Fruit set | Seed set |  |
| *Melochia villosa* | High | 0.02 | 0.81 | - | 0.36 | 7.14 | - | - | 23 |
|  | Low | 0.15 | 0.91 | - | 0.67 | 1.18 | - | - |  |
|  | Average | 0.08 | 0.86 | 0.17 | 0.52 | 4.16 | - | - |  |
| *Menyanthes trifoliata* | High | 0.05 | 0.88 | - | 0.06 | 11.86 | - | - | 27 |
|  | Low | 0.11 | 0.90 | - | 0.22 | 4.06 | - | - |  |
|  | Average | 0.08 | 0.89 | 0.49 | 0.14 | 7.96 | - | - |  |
| *Morinda pubescens* | High | 0.15 | 0.91 | - | 0.29 | 29.06 | - | - | This study |
|  | Low | 0.16 | 0.86 | - | 0.24 | 20.17 | - | - |  |
|  | Average | 0.15 | 0.89 | 0.54 | 0.26 | 24.62 | - | - |  |
| *Mussaenda decipiens* | High | 0.04 | 0.91 | - | 0.04 | 21.86 | - | - | 28 |
|  | Low | 0.26 | 0.96 | - | 0.30 | 2.74 | - | - |  |
|  | Average | 0.15 | 0.93 | 0.81 | 0.17 | 12.30 | - | - |  |
| *Mussaenda divaricata* | High | 0.25 | 0.97 | - | 0.32 | 2.25 | - | - | 28 |
|  | Low | 0.00 | 0.77 | - | 0.00 | 45.67 | - | - |  |
|  | Average | 0.13 | 0.87 | 0.84 | 0.16 | 23.96 | - | - |  |
| *Mussaenda erosa* | High | 0.01 | 0.91 | - | 0.00 | 16.11 | - | - | 28 |
|  | Low | 0.00 | 0.84 | - | 0.00 | 15.20 | - | - |  |
|  | Average | 0.01 | 0.88 | 0.74 | 0.00 | 15.65 | - | - |  |
| *Mussaenda hainanensis* | High | 0.00 | 0.84 | - | 0.00 | 61.86 | - | - | 28 |
|  | Low | 0.00 | 0.64 | - | 0.00 | 63.99 | - | - |  |
|  | Average | 0.00 | 0.74 | 0.45 | 0.00 | 62.92 | - | - |  |

Table S1 continued.
| Species | Level | Reciprocity index values |  |  |  |  | Reproductive success |  | Reference |
| --- | --- | --- | --- | --- | --- | --- | --- | --- | --- |
|  |  | COC | R & K | SAN | L & B | ARM | Fruit set | Seed set |  |
| *Mussaenda hirsutula* | High | 0.04 | 0.91 | - | 0.00 | 16.09 | - | - | 28 |
|  | Low | 0.04 | 0.88 | - | 0.06 | 14.92 | - | - |  |
|  | Average | 0.04 | 0.90 | 0.77 | 0.03 | 15.50 | - | - |  |
| *Mussaenda kwangsiensis* | High | 0.09 | 0.91 | - | 0.17 | 22.93 | - | - | 28 |
|  | Low | 0.00 | 0.67 | - | 0.00 | 44.93 | - | - |  |
|  | Average | 0.05 | 0.79 | 0.22 | 0.08 | 33.93 | - | - |  |
| *Mussaenda kwangtungensis* | High | 0.13 | 0.94 | - | 0.12 | 13.25 | - | - | 28 |
|  | Low | 0.00 | 0.56 | - | 0.00 | 121.79 | - | - |  |
|  | Average | 0.07 | 0.75 | 0.51 | 0.06 | 67.52 | - | - |  |
| *Mussaenda lancipetala* | High | 0.10 | 0.93 | - | 0.15 | 17.01 | - | - | 28 |
|  | Low | 0.19 | 0.94 | - | 0.37 | 4.29 | - | - |  |
|  | Average | 0.15 | 0.93 | 0.80 | 0.26 | 10.65 | - | - |  |
| *Mussaenda macrophylla* | High | 0.07 | 0.92 | - | 0.24 | 13.32 | - | - | 28 |
|  | Low | 0.16 | 0.93 | - | 0.20 | 5.10 | - | - |  |
|  | Average | 0.11 | 0.93 | 0.81 | 0.22 | 9.21 | - | - |  |
| *Mussaenda macrophylla* | High | 0.17 | 0.95 | - | 0.24 | 7.83 | - | - | 28 |
|  | Low | 0.01 | 0.84 | - | 0.00 | 22.99 | - | - |  |
|  | Average | 0.09 | 0.89 | 0.68 | 0.12 | 15.41 | - | - |  |
| *Mussaenda mollissima* | High | 0.09 | 0.92 | - | 0.07 | 29.18 | - | - | 28 |
|  | Low | 0.07 | 0.87 | - | 0.14 | 29.14 | - | - |  |
|  | Average | 0.08 | 0.89 | 0.58 | 0.10 | 29.16 | - | - |  |

Table S1 continued.
| Species | Level | Reciprocity index values |  |  |  |  | Reproductive success |  | Reference |
| --- | --- | --- | --- | --- | --- | --- | --- | --- | --- |
|  |  | COC | R & K | SAN | L & B | ARM | Fruit set | Seed set |  |
| *Mussaenda multinervis* | High | 0.00 | 0.83 | - | 0.00 | 61.44 | - | - | 28 |
|  | Low | 0.08 | 0.93 | - | 0.12 | 6.31 | - | - |  |
|  | Average | 0.04 | 0.88 | 0.65 | 0.06 | 33.87 | - | - |  |
| *Mussaenda parviflora* | High | 0.00 | 0.80 | - | 0.00 | 20.32 | - | - | 29 |
|  | Low | 0.14 | 0.92 | - | 0.47 | 1.77 | - | - |  |
|  | Average | 0.07 | 0.86 | 0.20 | 0.23 | 11.04 | - | - |  |
| *Mussaenda parviflora* | High | 0.02 | 0.83 | - | 0.06 | 19.60 | - | - | 29 |
|  | Low | 0.11 | 0.90 | - | 0.12 | 2.87 | - | - |  |
|  | Average | 0.06 | 0.87 | 0.39 | 0.09 | 11.24 | - | - |  |
| *Mussaenda pingbianensis* | High | 0.21 | 0.97 | - | 0.16 | 4.56 | - | - | 28 |
|  | Low | 0.00 | 0.54 | - | 0.00 | 132.09 | - | - |  |
|  | Average | 0.10 | 0.75 | 0.56 | 0.08 | 68.32 | - | - |  |
| *Mussaenda pubescens* | High | 0.16 | 0.95 | - | 0.40 | 2.56 | - | - | 28 |
|  | Low | 0.00 | 0.72 | - | 0.00 | 13.08 | - | - |  |
|  | Average | 0.08 | 0.84 | 0.53 | 0.20 | 7.82 | - | - |  |
| *Mussaenda pubescens* | High | 0.06 | 0.95 | - | 0.04 | 3.23 | - | - | 28 |
|  | Low | 0.00 | 0.71 | - | 0.00 | 22.78 | - | - |  |
|  | Average | 0.03 | 0.83 | 0.72 | 0.02 | 13.00 | - | - |  |
| *Mussaenda recurvata* | High | 0.03 | 0.87 | - | 0.00 | 34.60 | - | - | 30 |
|  | Low | 0.04 | 0.82 | - | 0.00 | 15.93 | - | - |  |
|  | Average | 0.04 | 0.84 | 0.47 | 0.00 | 25.26 | - | - |  |

Table S1 continued.
| Species | Level | Reciprocity index values |  |  |  |  | Reproductive success |  | Reference |
| --- | --- | --- | --- | --- | --- | --- | --- | --- | --- |
|  |  | COC | R & K | SAN | L & B | ARM | Fruit set | Seed set |  |
| *Nivenia argentea* | High | 0.04 | 0.93 | - | 0.25 | 17.21 | - | - | 31 |
|  | Low | 0.01 | 0.87 | - | 0.07 | 28.90 | - | - |  |
|  | Average | 0.02 | 0.90 | 0.68 | 0.16 | 23.06 | - | - |  |
| *Nivenia corymbosa* | High | 0.07 | 0.94 | - | 0.46 | 7.93 | - | - | 31 |
|  | Low | 0.08 | 0.92 | - | 0.23 | 4.58 | - | - |  |
|  | Average | 0.08 | 0.93 | 0.83 | 0.34 | 6.26 | - | - |  |
| *Nivenia inaequalis* | High | 0.02 | 0.93 | - | 0.42 | 40.78 | - | - | 31 |
|  | Low | 0.03 | 0.93 | - | 0.48 | 34.09 | - | - |  |
|  | Average | 0.02 | 0.93 | 0.81 | 0.45 | 37.44 | - | - |  |
| *Nymphoides montana* | High | 0.03 | 0.92 | - | 0.05 | 2.94 | 79.00 | 70.00 | 32 |
|  | Low | 0.24 | 0.96 | - | 0.40 | 0.37 | 84.00 | 83.00 |  |
|  | Average | 0.13 | 0.94 | 0.86 | 0.23 | 1.66 | - | - |  |
| *Nymphoides montana* | High | 0.01 | 0.89 | - | 0.05 | 4.87 | - | - | 32 |
|  | Low | 0.23 | 0.96 | - | 0.40 | 0.45 | - | - |  |
|  | Average | 0.12 | 0.92 | 0.77 | 0.23 | 2.66 | - | - |  |
| *Nymphoides montana* | High | 0.00 | 0.84 | - | 0.00 | 7.64 | 70.00 | 75.00 | 32 |
|  | Low | 0.23 | 0.95 | - | 0.35 | 0.42 | 79.00 | 74.00 |  |
|  | Average | 0.12 | 0.90 | 0.61 | 0.18 | 4.03 | - | - |  |
| *Ophiorrhiza japonica* | High | 0.17 | 0.96 | - | 0.46 | 1.90 | - | - | 33 |
|  | Low | 0.17 | 0.93 | - | 0.33 | 2.04 | - | - |  |
|  | Average | 0.17 | 0.94 | 0.89 | 0.39 | 1.97 | - | - |  |

Table S1 continued.
| Species | Level | Reciprocity index values |  |  |  |  | Reproductive success |  | Reference |
| --- | --- | --- | --- | --- | --- | --- | --- | --- | --- |
|  |  | COC | R & K | SAN | L & B | ARM | Fruit set | Seed set |  |
| *Oplonia nannophylla* | High | 0.10 | 0.94 | - | 0.50 | 3.20 | - | - | 34 |
|  | Low | 0.13 | 0.93 | - | 0.39 | 1.42 | - | - |  |
|  | Average | 0.12 | 0.93 | 0.82 | 0.44 | 2.31 | - | - |  |
| *Oplonia nannophylla* | High | 0.07 | 0.93 | - | 0.61 | 3.69 | - | - | 34 |
|  | Low | 0.10 | 0.91 | - | 0.42 | 2.68 | - | - |  |
|  | Average | 0.08 | 0.92 | 0.75 | 0.51 | 3.19 | - | - |  |
| *Palicourea coriacea* | High | 0.12 | 0.93 | - | 0.39 | 3.93 | - | - | 35 |
|  | Low | 0.11 | 0.91 | - | 0.49 | 4.38 | - | - |  |
|  | Average | 0.11 | 0.92 | 0.76 | 0.44 | 4.15 | - | - |  |
| *Palicourea coriacea* | High | 0.11 | 0.93 | - | 0.06 | 3.54 | - | - | 35 |
|  | Low | 0.15 | 0.92 | - | 0.40 | 4.24 | - | - |  |
|  | Average | 0.13 | 0.93 | 0.76 | 0.23 | 3.89 | - | - |  |
| *Palicourea coriacea* | High | 0.11 | 0.94 | - | 0.58 | 3.30 | - | - | 35 |
|  | Low | 0.14 | 0.94 | - | 0.33 | 2.18 | - | - |  |
|  | Average | 0.13 | 0.94 | 0.84 | 0.46 | 2.74 | - | - |  |
| *Palicourea coriacea* | High | 0.11 | 0.94 | - | 0.33 | 3.23 | - | - | 35 |
|  | Low | 0.16 | 0.94 | - | 0.32 | 1.84 | - | - |  |
|  | Average | 0.14 | 0.94 | 0.86 | 0.33 | 2.54 | - | - |  |
| *Palicourea coriacea* | High | 0.11 | 0.94 | - | 0.52 | 3.27 | - | - | 35 |
|  | Low | 0.14 | 0.94 | - | 0.33 | 2.47 | - | - |  |
|  | Average | 0.13 | 0.94 | 0.83 | 0.42 | 2.87 | - | - |  |

Table S1 continued.
| Species | Level | Reciprocity index values |  |  |  |  | Reproductive success |  | Reference |
| --- | --- | --- | --- | --- | --- | --- | --- | --- | --- |
|  |  | COC | R & K | SAN | L & B | ARM | Fruit set | Seed set |  |
| *Palicourea coriacea* | High | 0.11 | 0.94 | - | 0.27 | 3.16 | - | - | 35 |
|  | Low | 0.13 | 0.93 | - | 0.29 | 3.28 | - | - |  |
|  | Average | 0.12 | 0.93 | 0.83 | 0.28 | 3.22 | - | - |  |
| *Palicourea coriacea* | High | 0.17 | 0.95 | - | 0.58 | 2.34 | - | - | 35 |
|  | Low | 0.13 | 0.93 | - | 0.00 | 2.07 | - | - |  |
|  | Average | 0.15 | 0.94 | 0.85 | 0.29 | 2.21 | - | - |  |
| *Palicourea coriacea* | High | 0.12 | 0.94 | - | 0.36 | 3.66 | - | - | 35 |
|  | Low | 0.11 | 0.92 | - | 0.47 | 4.18 | - | - |  |
|  | Average | 0.12 | 0.93 | 0.77 | 0.41 | 3.92 | - | - |  |
| *Palicourea coriacea* | High | 0.10 | 0.94 | - | 0.25 | 3.04 | - | - | 35 |
|  | Low | 0.14 | 0.94 | - | 0.36 | 2.44 | - | - |  |
|  | Average | 0.12 | 0.94 | 0.84 | 0.31 | 2.74 | - | - |  |
| *Palicourea coriacea* | High | 0.12 | 0.94 | - | 0.27 | 3.21 | - | - | 35 |
|  | Low | 0.14 | 0.93 | - | 0.24 | 3.31 | - | - |  |
|  | Average | 0.13 | 0.94 | 0.83 | 0.25 | 3.26 | - | - |  |
| *Palicourea coriacea* | High | 0.15 | 0.94 | - | 0.31 | 3.26 | - | - | 35 |
|  | Low | 0.14 | 0.94 | - | 0.17 | 1.87 | - | - |  |
|  | Average | 0.15 | 0.94 | 0.83 | 0.24 | 2.57 | - | - |  |
| *Palicourea croceoides* | High | 0.03 | 0.91 | - | 0.05 | 7.00 | 54.90 | - | 36 |
|  | Low | 0.15 | 0.93 | - | 0.38 | 3.71 | 69.60 | - |  |
|  | Average | 0.09 | 0.92 | 0.72 | 0.22 | 5.35 | - | - |  |

Table S1 continued.
| Species | Level | Reciprocity index values |  |  |  |  | Reproductive success |  | Reference |
| --- | --- | --- | --- | --- | --- | --- | --- | --- | --- |
|  |  | COC | R & K | SAN | L & B | ARM | Fruit set | Seed set |  |
| *Palicourea croceoides* | High | 0.10 | 0.93 | - | 0.24 | 6.67 | 95.00 | - | 6 |
|  | Low | 0.09 | 0.88 | - | 0.36 | 9.13 | 81.25 | - |  |
|  | Average | 0.09 | 0.90 | 0.67 | 0.30 | 7.90 | - | - |  |
| *Palicourea padifolia* | High | 0.21 | 0.96 | - | 0.57 | 2.14 | 49.30 | 36.90 | 37 |
|  | Low | 0.00 | 0.76 | - | 0.00 | 17.75 | 40.50 | 29.70 |  |
|  | Average | 0.10 | 0.86 | 0.65 | 0.29 | 9.94 | - | - |  |
| *Palicourea rigida* | High | 0.03 | 0.96 | - | 0.32 | 0.04 | 56.60 | - | 38 |
|  | Low | 0.02 | 0.95 | - | 0.19 | 0.03 | 52.55 | - |  |
|  | Average | 0.02 | 0.96 | 0.89 | 0.25 | 0.03 | - | - |  |
| *Palicourea rigida* | High | 0.01 | 0.95 | - | 0.15 | 0.06 | 31.80 | - | 38 |
|  | Low | 0.03 | 0.94 | - | 0.26 | 0.05 | 41.70 | - |  |
|  | Average | 0.02 | 0.94 | 0.85 | 0.21 | 0.06 | - | - |  |
| *Palicourea tetragona* | High | 0.21 | 0.96 | - | 0.00 | 22.78 | - | - | 39 |
|  | Low | 0.16 | 0.95 | - | 0.20 | 31.97 | - | - |  |
|  | Average | 0.19 | 0.95 | 0.92 | 0.10 | 27.37 | - | - |  |
| *Pentanisia angustifolia* | High | 0.09 | 0.92 | - | 0.17 | 16.10 | 28.50 | - | 40 |
|  | Low | 0.11 | 0.91 | - | 0.25 | 11.86 | 33.10 | - |  |
|  | Average | 0.10 | 0.92 | 0.73 | 0.21 | 13.98 | - | - |  |
| *Pentanisia prunelloides* | High | 0.11 | 0.93 | - | 0.29 | 10.86 | 61.30 | - | 40 |
|  | Low | 0.14 | 0.93 | - | 0.12 | 5.89 | 61.30 | - |  |
|  | Average | 0.12 | 0.93 | 0.80 | 0.20 | 8.37 | - | - |  |

Table S1 continued.
| Species | Level | Reciprocity index values |  |  |  |  | Reproductive success |  | Reference |
| --- | --- | --- | --- | --- | --- | --- | --- | --- | --- |
|  |  | COC | R & K | SAN | L & B | ARM | Fruit set | Seed set |  |
| *Persicaria wugongshanensis* | High | 0.02 | 0.95 | - | 0.00 | 0.19 | - | - | 41 |
|  | Low | 0.50 | 0.98 | - | 0.85 | 0.01 | - | - |  |
|  | Average | 0.26 | 0.97 | 0.97 | 0.43 | 0.10 | - | - |  |
| *Plumbago auriculata* | High | 0.06 | 0.95 | - | 0.07 | 1.81 | 26.72 | - | 42 |
|  | Low | 0.24 | 0.96 | - | 0.42 | 0.78 | 23.11 | - |  |
|  | Average | 0.15 | 0.95 | 0.92 | 0.24 | 1.29 | - | - |  |
| *Polygonum hastatosagittatum* | High | 0.25 | 0.97 | - | 0.54 | 0.06 | - | 78.20 | 43 |
|  | Low | 0.11 | 0.92 | - | 0.17 | 0.12 | - | 74.10 |  |
|  | Average | 0.18 | 0.94 | 0.93 | 0.36 | 0.09 | - | - |  |
| *Polygonum jucundum* | High | 0.22 | 0.97 | - | 0.45 | 0.07 | - | - | 44 |
|  | Low | 0.00 | 0.83 | - | 0.00 | 0.40 | - | - |  |
|  | Average | 0.11 | 0.90 | 0.82 | 0.23 | 0.24 | - | - |  |
| *Primula allionii* | High | 0.08 | 0.91 | - | 0.44 | 3.70 | - | - | 45 |
|  | Low | 0.18 | 0.88 | - | 0.63 | 0.82 | - | - |  |
|  | Average | 0.13 | 0.89 | 0.58 | 0.53 | 2.26 | - | - |  |
| *Primula chungensis* | High | 0.20 | 0.96 | - | 0.71 | 1.22 | - | - | 46 |
|  | Low | 0.07 | 0.84 | - | 0.36 | 9.62 | - | - |  |
|  | Average | 0.14 | 0.90 | 0.63 | 0.53 | 5.42 | - | - |  |
| *Primula marginata* | High | 0.07 | 0.90 | - | 0.37 | 4.83 | - | - | 47 |
|  | Low | 0.18 | 0.89 | - | 0.60 | 0.87 | - | - |  |
|  | Average | 0.13 | 0.90 | 0.53 | 0.49 | 2.85 | - | - |  |

Table S1 continued.
| Species | Level | Reciprocity index values |  |  |  |  | Reproductive success |  | Reference |
| --- | --- | --- | --- | --- | --- | --- | --- | --- | --- |
|  |  | COC | R & K | SAN | L & B | ARM | Fruit set | Seed set |  |
| *Primula merrilliana* | High | 0.22 | 0.97 | - | 0.60 | 0.38 | - | 60.50 | 48 |
|  | Low | 0.19 | 0.95 | - | 0.30 | 0.28 | - | 70.70 |  |
|  | Average | 0.21 | 0.96 | 0.93 | 0.45 | 0.33 | - | - |  |
| *Primula mistassinica* | High | 0.20 | 0.94 | - | 0.56 | 0.60 | - | - | 49 |
|  | Low | 0.29 | 0.95 | - | 0.79 | 0.25 | - | - |  |
|  | Average | 0.25 | 0.94 | 0.86 | 0.68 | 0.43 | - | - |  |
| *Primula oreodoxa* | High | 0.16 | 0.95 | - | 0.76 | 1.32 | 90.10 | - | 50 |
|  | Low | 0.23 | 0.95 | - | 0.75 | 0.60 | 88.60 | - |  |
|  | Average | 0.19 | 0.95 | 0.89 | 0.75 | 0.96 | - | - |  |
| *Primula veris* | High | 0.05 | 0.89 | - | 0.20 | 12.97 | - | 57.14 | 51 |
|  | Low | 0.11 | 0.89 | - | 0.27 | 5.90 | - | 66.37 |  |
|  | Average | 0.08 | 0.89 | 0.57 | 0.24 | 9.44 | - | - |  |
| *Primula veris* | High | 0.07 | 0.91 | - | 0.20 | 8.55 | - | - | 51 |
|  | Low | 0.11 | 0.92 | - | 0.31 | 3.31 | - | - |  |
|  | Average | 0.09 | 0.92 | 0.75 | 0.26 | 5.93 | - | - |  |
| *Primula veris* | High | 0.08 | 0.92 | - | 0.30 | 9.25 | - | - | 51 |
|  | Low | 0.09 | 0.90 | - | 0.22 | 4.26 | - | - |  |
|  | Average | 0.08 | 0.91 | 0.72 | 0.26 | 6.76 | - | - |  |
| *Primula veris* | High | 0.05 | 0.90 | - | 0.10 | 11.20 | - | - | 51 |
|  | Low | 0.04 | 0.88 | - | 0.15 | 6.20 | - | - |  |
|  | Average | 0.04 | 0.89 | 0.67 | 0.12 | 8.70 | - | - |  |

Table S1 continued.
| Species | Level | Reciprocity index values |  |  |  |  | Reproductive success |  | Reference |
| --- | --- | --- | --- | --- | --- | --- | --- | --- | --- |
|  |  | COC | R & K | SAN | L & B | ARM | Fruit set | Seed set |  |
| *Primula veris* | High | 0.09 | 0.91 | - | 0.28 | 9.75 | - | - | 51 |
|  | Low | 0.13 | 0.91 | - | 0.31 | 4.20 | - | - |  |
|  | Average | 0.11 | 0.91 | 0.71 | 0.30 | 6.98 | - | - |  |
| *Primula veris* | High | 0.17 | 0.95 | - | 0.60 | 3.29 | - | 69.35 | 51 |
|  | Low | 0.16 | 0.93 | - | 0.32 | 2.24 | - | 67.86 |  |
|  | Average | 0.17 | 0.94 | 0.88 | 0.46 | 2.76 | - | - |  |
| *Primula veris* | High | 0.18 | 0.96 | - | 0.41 | 1.85 | - | - | 51 |
|  | Low | 0.23 | 0.95 | - | 0.48 | 1.23 | - | - |  |
|  | Average | 0.21 | 0.95 | 0.92 | 0.44 | 1.54 | - | - |  |
| *Primula veris* | High | 0.16 | 0.95 | - | 0.26 | 3.05 | - | - | 51 |
|  | Low | 0.11 | 0.92 | - | 0.24 | 2.97 | - | - |  |
|  | Average | 0.14 | 0.93 | 0.87 | 0.25 | 3.01 | - | - |  |
| *Primula veris* | High | 0.21 | 0.96 | - | 0.49 | 1.62 | - | - | 51 |
|  | Low | 0.16 | 0.94 | - | 0.19 | 1.78 | - | - |  |
|  | Average | 0.18 | 0.95 | 0.92 | 0.34 | 1.70 | - | - |  |
| *Primula veris* | High | 0.11 | 0.93 | - | 0.17 | 6.17 | - | - | 16 |
|  | Low | 0.14 | 0.89 | - | 0.26 | 4.80 | - | - |  |
|  | Average | 0.13 | 0.91 | 0.72 | 0.22 | 5.49 | - | - |  |
| *Psychotria asiatica* | High | 0.18 | 0.93 | - | 0.75 | 0.29 | 14.10 | - | 52 |
|  | Low | 0.11 | 0.85 | - | 0.29 | 0.57 | 0.00 | - |  |
|  | Average | 0.14 | 0.89 | 0.63 | 0.52 | 0.43 | - | - |  |

Table S1 continued.
| Species | Level | Reciprocity index values |  |  |  |  | Reproductive success |  | Reference |
| --- | --- | --- | --- | --- | --- | --- | --- | --- | --- |
|  |  | COC | R & K | SAN | L & B | ARM | Fruit set | Seed set |  |
| *Psychotria asiatica* | High | 0.24 | 0.95 | - | 0.73 | 0.22 | 15.40 | - | 52 |
|  | Low | 0.11 | 0.88 | - | 0.28 | 0.49 | 0.00 | - |  |
|  | Average | 0.18 | 0.91 | 0.74 | 0.50 | 0.36 | - | - |  |
| *Psychotria asiatica* | High | 0.21 | 0.94 | - | 0.77 | 0.22 | - | - | 52 |
|  | Low | 0.03 | 0.78 | - | 0.06 | 1.12 | - | - |  |
|  | Average | 0.12 | 0.86 | 0.51 | 0.42 | 0.67 | - | - |  |
| *Psychotria boninensis* | High | 0.16 | 0.93 | - | 0.60 | 0.48 | - | - | 53 |
|  | Low | 0.15 | 0.89 | - | 0.26 | 0.34 | - | - |  |
|  | Average | 0.15 | 0.91 | 0.71 | 0.43 | 0.41 | - | - |  |
| *Psychotria capitata* | High | 0.11 | 0.95 | - | 0.46 | 2.16 | 53.10 | - | 36 |
|  | Low | 0.12 | 0.93 | - | 0.37 | 2.22 | 59.00 | - |  |
|  | Average | 0.11 | 0.94 | 0.86 | 0.41 | 2.19 | - | - |  |
| *Psychotria capitata* | High | 0.10 | 0.94 | - | 0.35 | 2.08 | 80.70 | - | 35 |
|  | Low | 0.22 | 0.94 | - | 0.41 | 0.57 | 80.00 | - |  |
|  | Average | 0.16 | 0.94 | 0.85 | 0.38 | 1.32 | - | - |  |
| *Psychotria carthagenensis* | High | 0.08 | 0.92 | - | 0.37 | 1.84 | 30.37 | - | 54 |
|  | Low | 0.12 | 0.93 | - | 0.58 | 0.67 | 24.77 | - |  |
|  | Average | 0.10 | 0.92 | 0.75 | 0.48 | 1.25 | - | - |  |
| *Psychotria carthagenensis* | High | 0.00 | 0.79 | - | 0.04 | 7.55 | 3.72 | - | 54 |
|  | Low | 0.10 | 0.93 | - | 0.55 | 0.64 | 9.77 | - |  |
|  | Average | 0.05 | 0.86 | 0.28 | 0.29 | 4.10 | - | - |  |

Table S1 continued.
| Species | Level | Reciprocity index values |  |  |  |  | Reproductive success |  | Reference |
| --- | --- | --- | --- | --- | --- | --- | --- | --- | --- |
|  |  | COC | R & K | SAN | L & B | ARM | Fruit set | Seed set |  |
| *Psychotria carthagenensis* | High | 0.11 | 0.93 | - | 0.61 | 1.36 | 4.72 | - | 54 |
|  | Low | 0.15 | 0.93 | - | 0.61 | 0.57 | 9.90 | - |  |
|  | Average | 0.13 | 0.93 | 0.78 | 0.61 | 0.96 | - | - |  |
| *Psychotria cephalophora* | High | 0.16 | 0.95 | - | 0.55 | 0.36 | - | - | 55 |
|  | Low | 0.10 | 0.89 | - | 0.25 | 0.63 | - | - |  |
|  | Average | 0.13 | 0.92 | 0.80 | 0.40 | 0.50 | - | - |  |
| *Psychotria colorata* | High | 0.11 | 0.95 | - | 0.55 | 5.18 | 73.00 | - | 36 |
|  | Low | 0.17 | 0.95 | - | 0.76 | 2.66 | 77.80 | - |  |
|  | Average | 0.14 | 0.95 | 0.87 | 0.65 | 3.92 | - | - |  |
| *Psychotria deflexa* | High | 0.07 | 0.90 | - | 0.38 | 1.59 | 12.50 | - | 35 |
|  | Low | 0.16 | 0.90 | - | 0.62 | 0.39 | 3.60 | - |  |
|  | Average | 0.12 | 0.90 | 0.58 | 0.50 | 0.99 | - | - |  |
| *Psychotria deflexa* | High | 0.11 | 0.93 | - | 0.50 | 1.10 | 25.00 | - | 8 |
|  | Low | 0.11 | 0.90 | - | 0.40 | 0.81 | 30.00 | - |  |
|  | Average | 0.11 | 0.91 | 0.74 | 0.45 | 0.96 | - | - |  |
| *Psychotria deflexa* | High | 0.01 | 0.86 | - | 0.03 | 6.76 | 12.20 | - | 56 |
|  | Low | 0.00 | 0.77 | - | 0.00 | 4.78 | 3.60 | - |  |
|  | Average | 0.00 | 0.81 | 0.38 | 0.01 | 5.77 | - | - |  |
| *Psychotria gracilentia* | High | 0.08 | 0.91 | - | 0.42 | 1.57 | - | - | 36 |
|  | Low | 0.04 | 0.79 | - | 0.30 | 3.33 | - | - |  |
|  | Average | 0.06 | 0.85 | 0.30 | 0.36 | 2.45 | - | - |  |

Table S1 continued.
| Species | Level | Reciprocity index values |  |  |  |  | Reproductive success |  | Reference |
| --- | --- | --- | --- | --- | --- | --- | --- | --- | --- |
|  |  | COC | R & K | SAN | L & B | ARM | Fruit set | Seed set |  |
| *Psychotria hoffmannseggiana* | High | 0.07 | 0.92 | - | 0.33 | 1.06 | 84.80 | - | 36 |
|  | Low | 0.07 | 0.90 | - | 0.30 | 0.83 | 86.00 | - |  |
|  | Average | 0.07 | 0.91 | 0.72 | 0.32 | 0.95 | - | - |  |
| *Psychotria hoffmannseggiana* | High | 0.12 | 0.93 | - | 0.51 | 0.65 | 80.40 | - | 35 |
|  | Low | 0.14 | 0.90 | - | 0.53 | 0.51 | 85.20 | - |  |
|  | Average | 0.13 | 0.91 | 0.71 | 0.52 | 0.58 | - | - |  |
| *Psychotria homalosperma* | High | 0.11 | 0.92 | - | 0.24 | 23.30 | - | - | 57 |
|  | Low | 0.03 | 0.80 | - | 0.03 | 42.54 | - | - |  |
|  | Average | 0.07 | 0.86 | 0.43 | 0.14 | 32.92 | - | - |  |
| *Psychotria homalosperma* | High | 0.09 | 0.88 | - | 0.20 | 43.45 | - | - | 57 |
|  | Low | 0.03 | 0.80 | - | 0.07 | 47.33 | - | - |  |
|  | Average | 0.06 | 0.84 | 0.17 | 0.14 | 45.39 | - | - |  |
| *Psychotria jasminoides* | High | 0.12 | 0.94 | - | 0.26 | 3.25 | - | - | 58 |
|  | Low | 0.16 | 0.93 | - | 0.34 | 2.48 | - | - |  |
|  | Average | 0.14 | 0.94 | 0.79 | 0.30 | 2.86 | - | - |  |
| *Psychotria leiocarpa* | High | 0.05 | 0.87 | - | 0.33 | 2.37 | 80.00 | - | 56 |
|  | Low | 0.07 | 0.81 | - | 0.22 | 0.95 | 80.10 | - |  |
|  | Average | 0.06 | 0.84 | 0.23 | 0.27 | 1.66 | - | - |  |
| *Psychotria nervosa* | High | 0.04 | 0.94 | - | 0.03 | 44.43 | 4.67 | - | 59 |
|  | Low | 0.27 | 0.97 | - | 0.34 | 5.71 | 4.40 | - |  |
|  | Average | 0.16 | 0.96 | 0.92 | 0.19 | 25.07 | - | - |  |

Table S1 continued.
| Species | Level | Reciprocity index values |  |  |  |  | Reproductive success |  | Reference |
| --- | --- | --- | --- | --- | --- | --- | --- | --- | --- |
|  |  | COC | R & K | SAN | L & B | ARM | Fruit set | Seed set |  |
| *Psychotria nervosa* | High | 0.10 | 0.97 | - | 0.17 | 16.59 | 4.89 | - | 59 |
|  | Low | 0.30 | 0.97 | - | 0.30 | 4.63 | 4.82 | - |  |
|  | Average | 0.20 | 0.97 | 0.96 | 0.23 | 10.61 | - | - |  |
| *Psychotria peoppigiana* | High | 0.02 | 0.88 | - | 0.00 | 14.24 | - | - | 39 |
|  | Low | 0.15 | 0.90 | - | 0.17 | 5.20 | - | - |  |
|  | Average | 0.08 | 0.89 | 0.58 | 0.08 | 9.72 | - | - |  |
| *Psychotria poeppigiana* | High | 0.13 | 0.94 | - | 0.36 | 9.77 | - | - | 10 |
|  | Low | 0.18 | 0.93 | - | 0.43 | 4.86 | - | - |  |
|  | Average | 0.16 | 0.93 | 0.82 | 0.39 | 7.31 | - | - |  |
| *Psychotria serpens* | High | 0.14 | 0.94 | - | 0.40 | 0.50 | - | - | 60 |
|  | Low | 0.20 | 0.92 | - | 0.38 | 0.19 | - | - |  |
|  | Average | 0.17 | 0.93 | 0.81 | 0.39 | 0.34 | - | - |  |
| *Psychotria serpens* | High | 0.20 | 0.96 | - | 0.46 | 0.20 | - | - | 60 |
|  | Low | 0.14 | 0.90 | - | 0.21 | 0.30 | - | - |  |
|  | Average | 0.17 | 0.93 | 0.88 | 0.34 | 0.25 | - | - |  |
| *Psychotria serpens* | High | 0.14 | 0.94 | - | 0.28 | 0.46 | 53.40 | - | 60 |
|  | Low | 0.17 | 0.91 | - | 0.46 | 0.29 | 40.20 | - |  |
|  | Average | 0.16 | 0.92 | 0.80 | 0.37 | 0.38 | - | - |  |
| *Psychotria suerrensis* | High | 0.14 | 0.93 | - | 0.29 | 9.22 | - | - | 10 |
|  | Low | 0.19 | 0.93 | - | 0.33 | 2.68 | - | - |  |
|  | Average | 0.17 | 0.93 | 0.78 | 0.31 | 5.95 | - | - |  |

Table S1 continued.
| Species | Level | Reciprocity index values |  |  |  |  | Reproductive success |  | Reference |
| --- | --- | --- | --- | --- | --- | --- | --- | --- | --- |
|  |  | COC | R & K | SAN | L & B | ARM | Fruit set | Seed set |  |
| *Psychotria suterella* | High | 0.16 | 0.95 | - | 0.39 | 4.48 | 46.10 | - | 61 |
|  | Low | 0.17 | 0.94 | - | 0.43 | 3.08 | 62.00 | - |  |
|  | Average | 0.17 | 0.95 | 0.88 | 0.41 | 3.78 | - | - |  |
| *Psychotria trichophora* | High | 0.11 | 0.94 | - | 0.40 | 3.39 | 31.40 | - | 35 |
|  | Low | 0.12 | 0.91 | - | 0.31 | 2.59 | 19.70 | - |  |
|  | Average | 0.11 | 0.92 | 0.79 | 0.35 | 2.99 | - | - |  |
| *Psychotria trichophora* | High | 0.18 | 0.96 | - | 0.52 | 1.41 | 33.20 | - | 56 |
|  | Low | 0.23 | 0.94 | - | 0.44 | 1.05 | 19.30 | - |  |
|  | Average | 0.20 | 0.95 | 0.90 | 0.48 | 1.23 | - | - |  |
| *Pulmonaria angustifolia* | High | 0.07 | 0.94 | - | 0.10 | 2.85 | - | - | 62 |
|  | Low | 0.23 | 0.95 | - | 0.50 | 0.54 | - | - |  |
|  | Average | 0.15 | 0.94 | 0.84 | 0.30 | 1.69 | - | - |  |
| *Pulmonaria angustifolia* | High | 0.13 | 0.95 | - | 0.40 | 1.56 | - | - | 62 |
|  | Low | 0.08 | 0.88 | - | 0.20 | 1.39 | - | - |  |
|  | Average | 0.11 | 0.91 | 0.79 | 0.30 | 1.48 | - | - |  |
| *Pulmonaria collina* | High | 0.09 | 0.94 | - | 0.10 | 1.93 | - | - | 62 |
|  | Low | 0.16 | 0.93 | - | 0.27 | 0.67 | - | - |  |
|  | Average | 0.13 | 0.94 | 0.86 | 0.19 | 1.30 | - | - |  |
| *Pulmonaria collina* | High | 0.05 | 0.91 | - | 0.00 | 7.09 | - | - | 62 |
|  | Low | 0.18 | 0.94 | - | 0.30 | 0.94 | - | - |  |
|  | Average | 0.12 | 0.92 | 0.69 | 0.15 | 4.01 | - | - |  |

Table S1 continued.
| Species | Level | Reciprocity index values |  |  |  |  | Reproductive success |  | Reference |
| --- | --- | --- | --- | --- | --- | --- | --- | --- | --- |
|  |  | COC | R & K | SAN | L & B | ARM | Fruit set | Seed set |  |
| *Pulmonaria longifolia* | High | 0.16 | 0.96 | - | 0.10 | 1.01 | - | - | 62 |
|  | Low | 0.17 | 0.93 | - | 0.20 | 0.81 | - | - |  |
|  | Average | 0.16 | 0.95 | 0.90 | 0.15 | 0.91 | - | - |  |
| *Pulmonaria longifolia* | High | 0.12 | 0.94 | - | 0.18 | 2.76 | - | - | 16 |
|  | Low | 0.20 | 0.93 | - | 0.45 | 1.10 | - | - |  |
|  | Average | 0.16 | 0.94 | 0.84 | 0.31 | 1.93 | - | - |  |
| *Pulmonaria mollis* | High | 0.08 | 0.93 | - | 0.10 | 3.48 | - | - | 62 |
|  | Low | 0.14 | 0.92 | - | 0.10 | 1.24 | - | - |  |
|  | Average | 0.11 | 0.93 | 0.77 | 0.10 | 2.36 | - | - |  |
| *Pulmonaria obscura* | High | 0.09 | 0.95 | - | 0.00 | 1.56 | - | - | 62 |
|  | Low | 0.19 | 0.93 | - | 0.20 | 0.69 | - | - |  |
|  | Average | 0.14 | 0.94 | 0.87 | 0.10 | 1.13 | - | - |  |
| *Pulmonaria obscura* | High | 0.04 | 0.92 | - | 0.10 | 3.40 | - | - | 62 |
|  | Low | 0.18 | 0.93 | - | 0.20 | 0.70 | - | - |  |
|  | Average | 0.11 | 0.93 | 0.80 | 0.15 | 2.05 | - | - |  |
| *Pulmonaria obscura* | High | 0.15 | 0.95 | - | 0.10 | 1.64 | - | - | 62 |
|  | Low | 0.16 | 0.92 | - | 0.10 | 0.85 | - | - |  |
|  | Average | 0.15 | 0.94 | 0.86 | 0.10 | 1.25 | - | - |  |
| *Pulmonaria officinalis* | High | 0.14 | 0.96 | - | 0.39 | 1.72 | - | 61.50 | 63 |
|  | Low | 0.03 | 0.90 | - | 0.00 | 1.82 | - | 59.30 |  |
|  | Average | 0.08 | 0.93 | 0.87 | 0.19 | 1.77 | - | - |  |

Table S1 continued.
| Species | Level | Reciprocity index values |  |  |  |  | Reproductive success |  | Reference |
| --- | --- | --- | --- | --- | --- | --- | --- | --- | --- |
|  |  | COC | R & K | SAN | L & B | ARM | Fruit set | Seed set |  |
| *Pulmonaria officinalis* | High | 0.08 | 0.94 | - | 0.11 | 3.11 | - | 61.50 | 63 |
|  | Low | 0.07 | 0.92 | - | 0.05 | 1.42 | - | 59.30 |  |
|  | Average | 0.07 | 0.93 | 0.83 | 0.08 | 2.26 | - | - |  |
| *Pulmonaria officinalis* | High | 0.07 | 0.92 | - | 0.14 | 5.44 | - | - | 62 |
|  | Low | 0.02 | 0.90 | - | 0.00 | 2.58 | - | - |  |
|  | Average | 0.05 | 0.91 | 0.70 | 0.07 | 4.01 | - | - |  |
| *Pulmonaria officinalis* | High | 0.07 | 0.92 | - | 0.10 | 4.17 | - | - | 62 |
|  | Low | 0.15 | 0.92 | - | 0.30 | 1.00 | - | - |  |
|  | Average | 0.11 | 0.92 | 0.73 | 0.20 | 2.58 | - | - |  |
| *Pulmonaria officinalis* | High | 0.12 | 0.96 | - | 0.21 | 1.76 | - | - | 62 |
|  | Low | 0.04 | 0.90 | - | 0.00 | 1.77 | - | - |  |
|  | Average | 0.08 | 0.93 | 0.87 | 0.11 | 1.76 | - | - |  |
| *Pulmonaria officinalis* | High | 0.10 | 0.94 | - | 0.10 | 3.10 | - | - | 62 |
|  | Low | 0.08 | 0.89 | - | 0.10 | 2.23 | - | - |  |
|  | Average | 0.09 | 0.91 | 0.79 | 0.10 | 2.66 | - | - |  |
| *Pulmonaria officinalis* | High | 0.08 | 0.92 | - | 0.10 | 4.29 | - | - | 62 |
|  | Low | 0.07 | 0.90 | - | 0.10 | 1.47 | - | - |  |
|  | Average | 0.07 | 0.91 | 0.73 | 0.10 | 2.88 | - | - |  |
| *Pulmonaria officinalis* | High | 0.14 | 0.95 | - | 0.47 | 1.82 | - | - | 64 |
|  | Low | 0.16 | 0.93 | - | 0.57 | 1.03 | - | - |  |
|  | Average | 0.15 | 0.94 | 0.85 | 0.52 | 1.42 | - | - |  |

Table S1 continued.
| Species | Level | Reciprocity index values |  |  |  |  | Reproductive success |  | Reference |
| --- | --- | --- | --- | --- | --- | --- | --- | --- | --- |
|  |  | COC | R & K | SAN | L & B | ARM | Fruit set | Seed set |  |
| *Pulmonaria saccharata* | High | 0.08 | 0.92 | - | 0.20 | 4.49 | - | - | 62 |
|  | Low | 0.17 | 0.93 | - | 0.10 | 0.98 | - | - |  |
|  | Average | 0.13 | 0.93 | 0.75 | 0.15 | 2.73 | - | - |  |
| *Pulmonaria saccharata* | High | 0.11 | 0.94 | - | 0.10 | 3.01 | - | - | 62 |
|  | Low | 0.14 | 0.91 | - | 0.20 | 1.93 | - | - |  |
|  | Average | 0.13 | 0.93 | 0.82 | 0.15 | 2.47 | - | - |  |
| *Salvia brandegeei* | High | 0.18 | 0.96 | - | 0.62 | 1.42 | - | 52.00 | 65, 66 |
|  | Low | 0.13 | 0.93 | - | 0.46 | 1.66 | - | 49.00 |  |
|  | Average | 0.16 | 0.94 | 0.88 | 0.54 | 1.54 | - | - |  |
| *Schizomussaenda henryi* | High | 0.06 | 0.93 | - | 0.20 | 23.53 | - | - | 28 |
|  | Low | 0.22 | 0.96 | - | 0.28 | 6.88 | - | - |  |
|  | Average | 0.14 | 0.94 | 0.86 | 0.24 | 15.20 | - | - |  |
| *Turnera scabra* | High | 0.10 | 0.93 | - | 0.21 | 4.36 | - | - | 67 |
|  | Low | 0.02 | 0.81 | - | 0.00 | 8.13 | - | - |  |
|  | Average | 0.06 | 0.87 | 0.64 | 0.11 | 6.24 | - | - |  |
| *Turnera subulata* | High | 0.21 | 0.96 | - | 0.52 | 1.48 | - | 33.72 | 67, 68 |
|  | Low | 0.20 | 0.92 | - | 0.34 | 1.48 | - | 33.47 |  |
|  | Average | 0.21 | 0.94 | 0.88 | 0.43 | 1.48 | - | - |  |
| *Turnera subulata* | High | 0.11 | 0.92 | - | 0.16 | 5.65 | 97.00 | 43.40 | 69 |
|  | Low | 0.17 | 0.91 | - | 0.19 | 2.35 | 87.00 | 71.95 |  |
|  | Average | 0.14 | 0.92 | 0.74 | 0.18 | 4.00 | - | - |  |

Table S1 continued.
| Species | Level | Reciprocity index values |  |  |  |  | Reproductive success |  | Reference |
| --- | --- | --- | --- | --- | --- | --- | --- | --- | --- |
|  |  | COC | R & K | SAN | L & B | ARM | Fruit set | Seed set |  |
| *Turnera ulmifolia* | High | 0.11 | 0.93 | - | 0.40 | 3.79 | - | - | 67 |
|  | Low | 0.07 | 0.88 | - | 0.10 | 3.64 | - | - |  |
|  | Average | 0.09 | 0.90 | 0.74 | 0.25 | 3.72 | - | - |  |
| *Vismia guianensis* | High | 0.64 | 0.98 | - | 1.00 | 0.00 | 23.00 | - | 70 |
|  | Low | 0.65 | 0.96 | - | 1.00 | 0.00 | 17.00 | - |  |
|  | Average | 0.65 | 0.97 | 0.97 | 1.00 | 0.00 | - | - |  |

